# Direct RNA Sequencing reveals epitranscriptomic regulation of brain cells and Alzheimer’s Disease pathology

**DOI:** 10.64898/2026.05.18.724443

**Authors:** Ashley Byrne, Jonathan Hoover, Jessica Lund, Ai Zhang, Xiwei Shan, Anup Dutt Sharma, Xuewen Chen, Ana Xavier-Magalhães, Lilian Phu, Meena Choi, Ahmet Kurdoglu, Christopher M Rose, Alessandro Ori, Yuxin Liang, Alicia Nugent, Claire Jeong, Zora Modrusan, William Stephenson, Daniel Le

## Abstract

Alternative mRNA splicing and post-transcriptional RNA modification are key mechanisms that regulate transcript function; however, their role in neuronal activity and neurodegenerative disease remains poorly defined. In this study, we evaluated two nanopore-based long-read sequencing (LR-seq) formats: cDNA-PCR sequencing (CPS) and direct RNA sequencing (DRS). We then applied DRS to profile both full-length isoforms and RNA modifications in major brain cell types derived from induced pluripotent stem cells (iPSCs) and post-mortem Alzheimer’s disease (AD) brains. Relative to CPS, DRS achieved higher accuracy and sensitivity for transcript quantification, *de novo* transcript model construction, and open reading frame (ORF) annotation across neuropathological gene sets. Focusing on iPSC-derived neurons, we built a multi-omic atlas to connect transcriptional output with translational engagement and protein abundance, by integrating DRS-based mRNA abundance, N^6^-methyladenosine (m6A) status and poly(A) tail length with ribosome profiling (Ribo-seq) and mass spectrometry (MS). The combination of DRS and Ribo-seq data demonstrated synergism in predicting protein abundance. This analysis also uncovered a significant inverse relationship between m6A modification and mRNA abundance, which was dependent on the engagement of the ribosomal A-site. Lastly, we applied DRS to the epitranscriptomic analysis of AD brain samples, demonstrating that m6A profiles can be used to distinguish early-versus late-stage disease.

## Introduction

The rapid evolution of RNA sequencing technologies has profoundly enhanced our understanding of cellular processes, enabling unprecedented insight into cellular functional complexity. A major breakthrough came with the emergence of LR-seq platforms, pioneered by Oxford Nanopore Technologies and Pacific Biosciences, which allow for high-resolution, comprehensive characterization of RNA transcripts ^1–5^. By facilitating the discovery of novel, disease-relevant transcripts that were previously undetected with short-read sequencing, LR-seq has opened new opportunities for diagnostic and therapeutic advancements ^6^. In recent years, nanopore-based DRS has further transformed the RNA analysis landscape ^7^. Unlike approaches that detect cDNA, which require both reverse transcription (RT) and PCR, DRS directly sequences native RNA molecules, circumventing biases inherent to RT-PCR. This attribute enables quantitative and unbiased measurements of RNA isoforms in addition to facilitating RNA base modification detection. These features make DRS particularly well-suited for capturing the intrinsic complexity and abundance of RNA molecules within cells.

One of the most promising applications of DRS lies in its ability to interrogate the epitranscriptome - the collection of chemical modifications on RNA molecules that modulate transcript function and cellular processes ^8,9^. These epitranscriptomic marks influence crucial mechanisms such as RNA stability, splicing, translation, and immune sensing. Furthermore, the colocalization of multiple RNA modifications may act as a regulatory “code,” analogous to the histone code in epigenomic regulation ^10^. To decipher this complex regulatory grammar, it is essential to accurately identify the types, positions, and stoichiometries of RNA modifications. DRS offers distinct advantages in this context, including single-molecule resolution, single-base precision, and the simultaneous detection of multiple modification types. Indeed, DRS has already been successfully applied to map colocalized RNA modifications, revealing previously uncharacterized aspects of transcript regulation ^11,12^.

Among the diverse array of RNA modifications, m6A is the most abundant modification in eukaryotic mRNAs ^13–16^. This modification plays critical roles in fundamental biological processes, with particular significance in brain development and function ^17–20^. Dysregulation of m6A-associated pathways has been linked to several neurological disorders, underscoring the importance of technologies like DRS to elucidate the precise role of m6A in the brain ^21–24^. The ability of DRS to directly detect and quantify m6A modifications provides an unprecedented opportunity to explore the dynamic interplay between m6A and neuronal processes, potentially advancing our understanding of brain health and disease. Furthermore, improvements to basecalling algorithms continue to expand DRS capabilities, enabling the detection of additional RNA modifications such as pseudouridine (PseU), inosine, 5-methylcytosine (m5C) and many others ^12,25–28^.

Here, we utilize DRS to characterize the transcriptomic and epitranscriptomic landscape of major brain cell types and of the frontal cortex from AD patients. Our findings demonstrate that DRS outperforms CPS in key performance metrics, including transcript quantification, *de novo* transcript model construction, and ORF detection. Validation *via* MS confirmed that ORFs identified using DRS are translated, with many implicated in neuronal processes. Furthermore, DRS enabled the detection of RNA modifications, such as m6A, within a neuronal model system. Notably, we observed a significant inverse relationship between m6A levels and transcript abundance, particularly at codons engaged with the ribosomal A-site. Lastly, both gene-level and single-nucleotide differential m6A analysis revealed global hypomethylation among late-stage AD samples. Together, these findings highlight DRS as a powerful and versatile technology for exploring the complex interplay between transcriptomic regulation, RNA modifications, and neurological disorders like AD.

## Results

### Section 1: DRS exhibits enhanced transcript quantitation accuracy and detection sensitivity

#### DRS demonstrates higher accuracy than CPS in quantifying synthetic control transcripts

To assess the quantitative performance of DRS and CPS, total RNA derived from SH-SY5Y neuroblastoma cells supplemented with External RNA Control Consortium (ERCC) and Spike-in RNA Variant (SIRV) control mixtures was used for benchmarking (**Figure 1a**). CPS libraries were multiplexed to approximate DRS flow cell read yield (DRS = 9.7-11.5 M; CPS = 33.8-41.1 M), then each sample was depth normalized to approximately 9.7 M reads. *IsoQuant* ^29^ was used to quantify control transcripts for both sequencing formats, revealing similar accuracy based on the Pearson correlation coefficient of observed ERCC transcript abundances relative to expected values: DRS = 0.94 and CPS = 0.93 (*p*-values < 0.05; **EDF 1a**). With respect to SIRVs, DRS demonstrated superior accuracy over CPS, achieving a lower root-mean-square error (RMSE) for isoform quantitation **(Figure 1b,c**).

**Figure 1.**
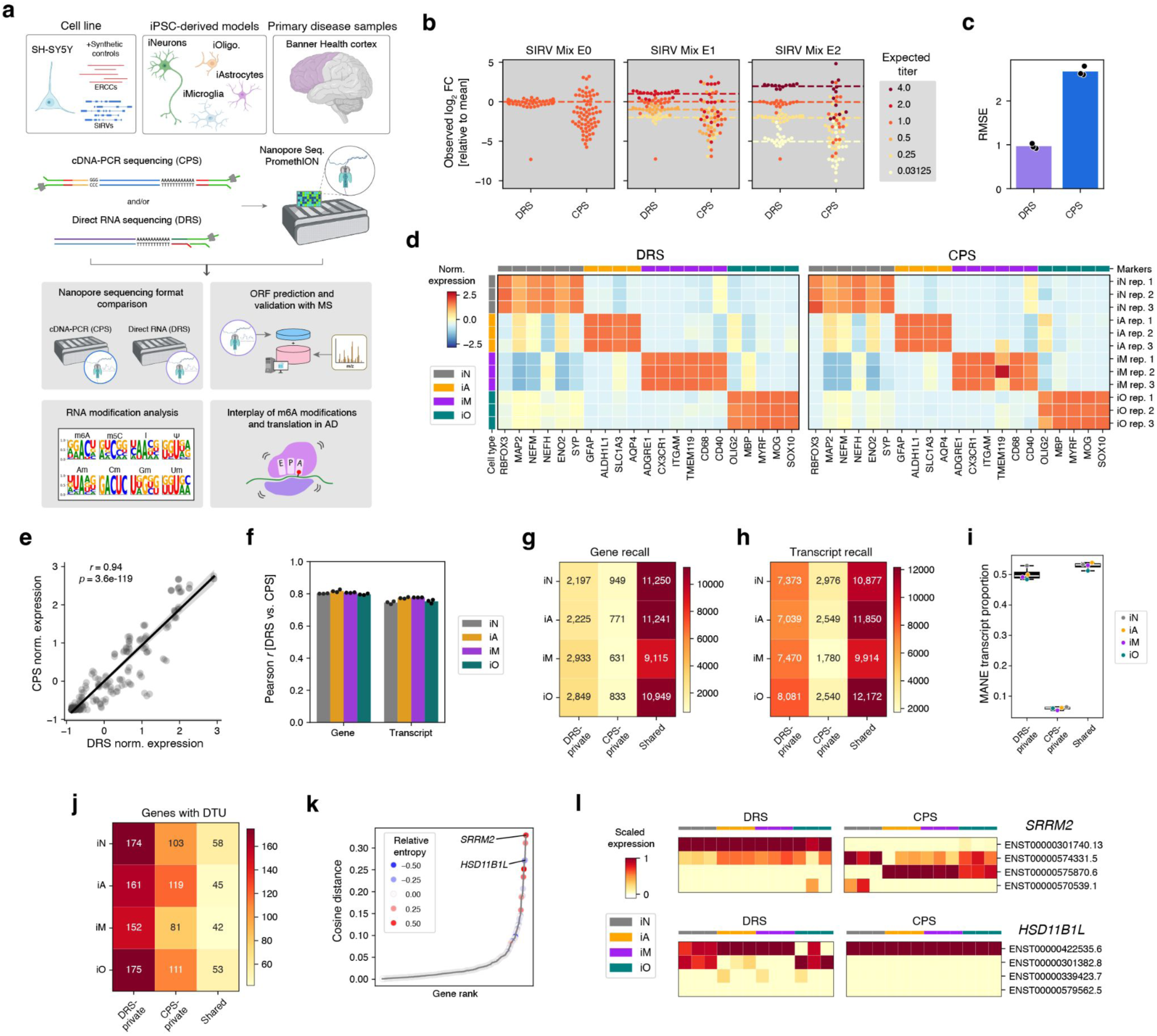
**Quantitation accuracy**. **a**) Schematic overview of samples, sequencing formats and analyses in this study. **b**) Swarmplots with each point representing a recalled transcript and displaying sequencing formats versus mean relative titer across replicates (n = 3): (left) Mix E0 = equimolar, (center) Mix E1 = narrow range, (right) Mix E2 = broad range. Color-coded horizontal dashed lines indicate expected concentrations, projected on a relative fold-change scale based on the mean TPM across all transcripts. **c**) Barplots depicting root mean squared error (RMSE) for each sequencing format across replicates (n = 3). **d**) Heatmaps displaying respective iPSC-derived cell type marker gene expression for DRS (left) and CPS (right), across replicates (n = 3). **e**) Scatterplot showing marker gene expression correlation between DRS and CPS. Pearson correlation coefficient and P value are shown, across replicates (n = 3). **f**) Barplot showing gene-and transcript-level Pearson correlation coefficients comparing DRS and CPS expression values, disaggregated by iPSC-derived cell type (n = 3). Genes and transcripts with CPM > 1. **g)** Heatmap displaying the number of recalled genes, found in all replicates (n = 3), unique to sequencing format (*i.e.* dRNA-and cDNA-private) in addition to those found in both formats (*i.e.* Shared) for each iPSC-derived cell type. **h**) Similar to before, except for transcript recall. **i**) Boxplot showing mean MANE label proportion, across replicates (n = 3), among recalled transcripts for each iPSC-derived cell type, separated into dRNA-private, cDNA-private and shared transcript recall groups. **j**) Heatmap displaying number of genes exhibiting differential transcript usage (DTU), across replicates (n = 3), separated by iPSC-derived cell type versus dRNA-private, cDNA-private and shared groups. **k**) Scatterplot showing genes with a DTU event ranked by cosine distance between DRS and CPS transcript expression profiles (y-axis). Each gene is represented by a point that is color-coded based on the relative entropy between DRS and CPS transcript expression profiles. **l**) (top) For the SRRM2 gene, heatmap showing normalized transcript expression for each replicate, grouped by iPSC-derived cell type and separated by sequencing format. The four transcripts with the highest mean expression are shown, across replicates (n = 3). (bottom) Similar to before, except for the HSD11B1L gene.

#### DRS yields longer reads than CPS but has a lower mapping rate

Expanding our analysis beyond SH-SY5Y cells, we examined four major brain cell types derived from iPSCs: iNeurons (iN), iAstrocytes (iA), iMicroglia (iM), and iOligodendrocytes (iO). Cell type identities were validated using marker gene expression ^30^ (**Figure 1d**), and showed strong agreement between DRS and CPS (Pearson *r* = 0.94; *p*-value = 3.6e-119; **Figure 1e**). We also observed good correspondence between DRS and CPS across all detected genes and transcripts, with mean gene-level correlation coefficient slightly higher than that of transcripts (mean Pearson r = 0.81 and 0.76, respectively; **Figure 1f**). Next, we compared the read length distributions between the two sequencing formats. DRS consistently produced longer reads, with a median length across replicates and cell types of 1,587 nt, compared to 903 nt for CPS (**EDF 1b**). DRS also exhibited cell type-specific variation in read length distributions, with peaks ranging between 878-1789 nt. In contrast, read length distribution peaks from CPS spanned a narrower range of 899-915 nt. The observed cell-type specific read length distributions indicate that DRS offers greater transcriptome sampling fidelity compared to CPS, which can be biased by RT-PCR.

Across the DRS libraries, we observed read yields between 8.6-19.2 M with a median utilization rate of 0.55, defined as the fraction of informative read counts used to determine gene expression (**EDF 1c**). In comparison, CPS libraries yielded between 15.8-27.3 M read with a markedly higher median utilization rate of 0.78 (**EDF 1d**). This discrepancy led us to investigate the unutilized portion of reads, composed of three subtypes: ambiguous (*i.e.* aligned to multiple transcripts), unmapped and intergenic (**EDF 1e**). While the proportion of ambiguous and intergenic reads was similar between sequencing formats, we observed a much higher unmapped fraction among DRS libraries (**EDF 1f**). In particular, the iN samples exhibited a median unmapped fraction of 0.47, about 2-fold higher than other DRS samples (range = 0.24-0.29). The iN samples exhibited a large subpopulation of relatively short reads, approximately 50-300 nt (**EDF 1b**), which led to the hypothesis that the elevated unmapped fraction is partly explained by low quality short reads. Comparing across all samples, both median read length and mean PRHED score of unmapped reads correlated with the overall mapping rate (Pearson r = 0.61 and 0.86, respectively; **EDF 1g**).

#### DRS exhibits higher detection sensitivity for genes, transcripts, and differential transcript usage (DTU) events

Compared to CPS, DRS showed substantially higher recall sensitivity across all iPSC-derived samples, identifying 3.2-and 3.0-fold more private annotated genes and transcripts, respectively (**Figure 1g,h**). Notably, the private transcripts identified by DRS included 8.1-fold more Matched Annotation from NCBI and EMBL-EBI (MANE) transcripts, which represent well-supported and conserved isoforms (**Figure 1i**).The proportion of MANE transcripts within DRS private transcripts (median = 0.50) was comparable to that of shared transcripts (median = 0.53); whereas, CPS private transcripts contained a significantly lower proportion of MANE isoforms (median = 0.06). These findings suggest that the relatively limited number of transcripts uniquely recalled by CPS do not reflect canonical isoforms, possibly due to sampling bias from abortive RT and PCR.

Given its higher recall sensitivity, DRS also identified more DTU events. For “query-versus-rest” permutation tests designed to identify reciprocal differential major isoform expression (*e.g.* iN versus iA + iM + iO), DRS consistently detected a greater number of private DTU events (**Figure 1j**). Interestingly, the overlap of DTU events between DRS and CPS was limited, consistent with substantial format-specific differences in transcript detection. To further characterize these differences, we calculated cosine distance and relative Shannon entropy between DRS and CPS isoform expression profiles per gene, revealing significant variation in the breadth of transcript utilization (**Figure 1k**). Such sequencing format discrepancies may dramatically impact biological interpretations. For example, DRS determined relatively high expression of the long transcript ENST00000301740.13 of SRRM2, a RNA splicing factor important in neurodevelopment ^31^; whereas, CPS reads were composed of shorter splice variants (**Figure 1l**, **EDF 1h**). Similarly, for the glucocorticoid-regulating enzyme HSD11B1L, DRS captured multiple long isoforms, while CPS primarily detected the short transcript ENST00000422535.6 (**EDF 1i**). These discrepancies underscore the influence of sequencing format on isoform expression profiles that underpin subsequent biological inferences.

### Section 2: DRS identifies unannotated protein-coding transcripts in iNeurons and AD

#### Transcript model characterization and validation with MS

Considering the observed disparity in annotated transcript detection between DRS and CPS, we hypothesized that the two sequencing formats might generate distinct sets of transcript models with protein-coding potential. Thus, we developed a workflow to identify reproducibly-detected predicted open reading frames (RD-ORFs) coupled with protein translation validation using MS. Transcript models were constructed using *IsoQuant* with ambiguous or inconsistent reads relative to GENCODE version 46 annotations (05/2024 release). Models with expression greater than one transcript per million (TPM) in all replicates were subsequently evaluated for protein-coding potential using *TD2* ^32^, based on the *PSAURON* tool ^33^. Transcripts with *PSAURON* scores exceeding 0.9 (indicating high protein-coding potential) were determined to be RD-ORFs. Notably, a larger proportion of reproducibly-detected transcript models from DRS displayed protein-coding potential, compared to CPS: median of 94% versus 88%, respectively (**Figure 2a**). Moreover, the median RD-ORF predicted polypeptide length from DRS was 2-fold longer than that from CPS (**Figure 2b**), which aligns with previous findings indicating that DRS is more capable of characterizing longer transcripts. Across all iPSC-derived samples, DRS identified an average of 1.8-fold more private RD-ORFs compared to CPS (**Figure 2c**). Moreover, the number of private RD-ORFs was consistently greater than the number found by both sequencing formats. This indicates a substantial degree of mutual exclusivity between DRS and CPS that can be exploited to discover a broader range of transcript species. Importantly, the number of ORFs detected by either DRS or CPS is not dependent on overall gene expression levels (mean Spearman ⍴ = 0.10; **EDF 2a**); rather, the detection of RD-ORFs seems to reflect differences in transcript utilization within the cell.

**Figure 2.**
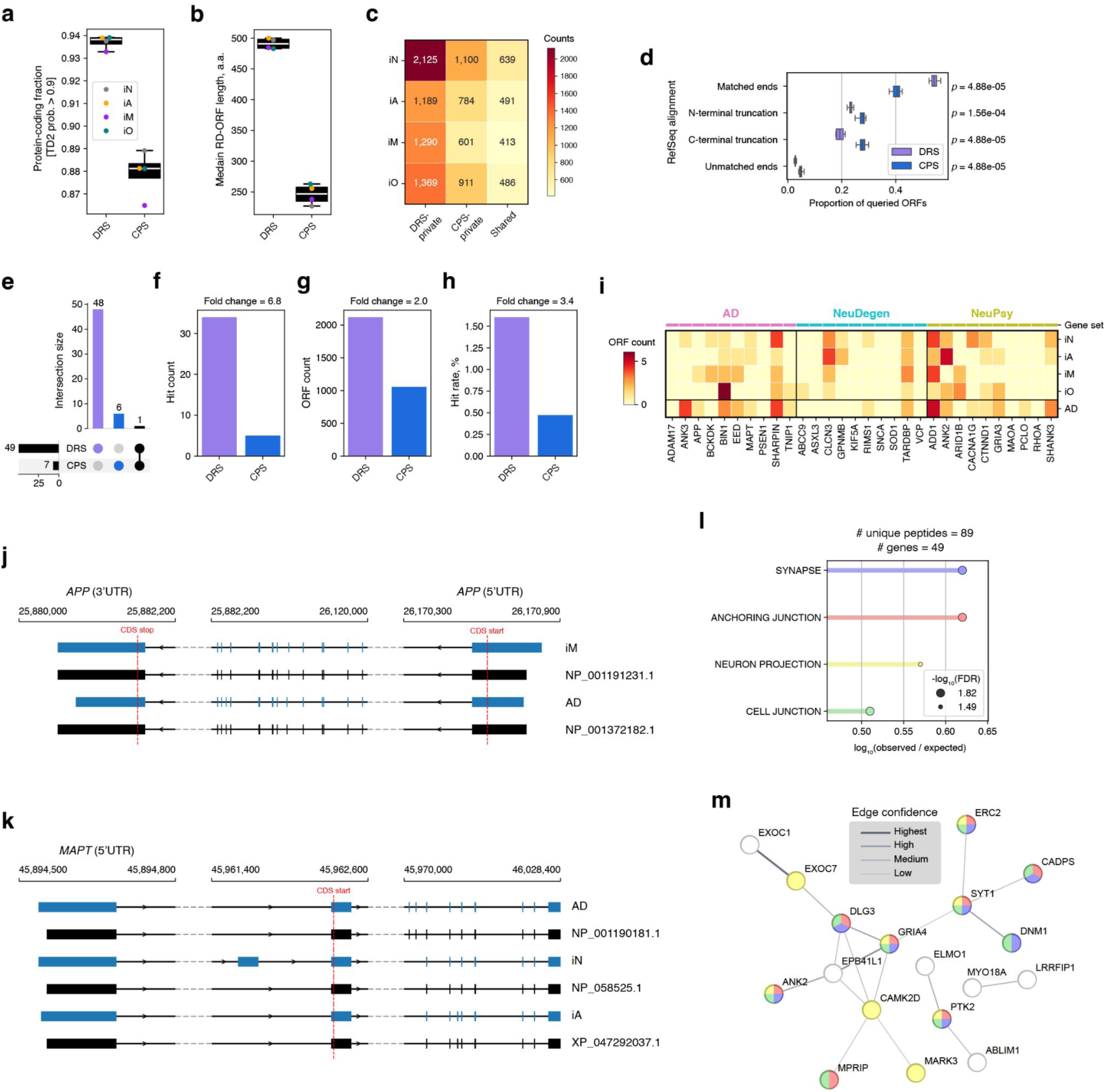
Putative ORF detection. **a**) Boxplot showing the fraction of passing RD-ORFs, across replicates (n = 3), using DRS versus CPS, disaggregated by iPSC-derived cell type. **b**) Boxplot similar to before, except displaying median RD-ORF lengths. **c**) Heatmap showing number of RD-ORFs, across replicates (n = 3), unique to a sequencing format or shared, disaggregated by iPSC-derived cell type. **d**) Boxplot showing the alignment classification proportion of queried ORFs against the RefSeq protein database. Two-sided Mann-Whitney *U* test FDR comparing DRS versus CPS shown along the right y-axis (n = 12). **e**) UpSet plot illustrating the distribution of peptide mass spectra identified using peptide databases derived from either DRS or CPS. **f-h**) Barplots comparing DRS versus CPS with respect to hit count (*i.e.* number of sequencing format-specific RD-ORFs with MS support), ORF count (*i.e.* number of supplemented polypeptide sequences) and hit rate (*i.e.* percentage of RD-ORFs with MS support relative to the total number of supplemented polypeptide sequences). **i**) Heatmap depicting the ORF counts per gene belonging to specific gene sets: Alzheimer’s disease (AD), neurodegeneration (NeuDegen) and neuropsychiatric (NeuPys). **j**) IGV gene tracks for *APP* transcript models zoomed into UTRs. **k**) IGV gene tracks for *MAPT* transcript models zoomed into the 5’-UTR. **l**) GSEA plot showing gene set terms that were significantly enriched with query genes with a detected novel peptide, derived from AD samples. Results were generated using the “Analysis” function at https://string-db.org/. The x-axis shows enrichment strength, log_10_(observed / expected), which is based on the observed number of query genes overlapping a gene set compared to a random query set. For each term, the FDR is depicted as circles of various diameters according to the negative log_10_ transform. **m**) Interaction network plot depicting connected query genes. Each node is a gene with edges indicating an interaction with another gene. Edge thickness indicates the interaction confidence score.

We then characterized the alterations in ORFs that distinguish them from annotated transcripts. DRS produced a greater proportion of transcript models with known intron retention and showed fewer alignment artifacts compared to CPS (two-sided Mann-Whitney *U* test FDR < 0.05; **EDF 2b**). A granular classification of alteration events revealed several structural changes that were more prevalent in DRS, such as exon skipping and exon gain, along with previously noted intron retention (two-sided Mann-Whitney *U* test FDR < 0.05; **EDF 2c**). In contrast, CPS frequently generated novel intron events: unannotated intron chains (alternative_structure), introns within an exon (extra_intron), and unclassified intron alteration (intron_alternation). Taken together, these sequencing format-specific transcript alterations lead to an expanded set of predicted polypeptide sequences, compared to those encoded by annotated transcripts.

Next, we queried the predicted polypeptide sequences against the RefSeq protein catalog (08/2024 release) using *BLASTp* ^34^. For each predicted peptide sequence, a top-ranked homologous reference protein was identified based on query sequence coverage and percent identity. DRS showed a 1.4-fold greater frequency of alignments where both the N-and C-termini matched the reference protein. In contrast, CPS exhibited higher rates of N-and C-terminal truncations and doubly unmatched ends (two-sided Mann-Whitney *U* test FDR < 0.05; **Figure 2d)**. Moreover, we observed a 1.3-fold increase in the number of gaps per alignment from DRS compared to CPS (two-sided Mann-Whitney *U* test FDR < 0.05; **EDF 2d)**. This finding is consistent with previously reported elevated levels of deletion errors associated with DRS ^35^. Intriguingly, both DRS and CPS display mean gap lengths around 13-16 nt, which may indicate alternative splicing of 3-27 nt micro exons predominantly found in neurons ^36^. Overall, most polypeptide sequences predicted from RD-ORFs showed good alignment with a reference protein, serving as external validation. The few sequences that did not align might be due to sequencing or library preparation artifacts, or they could represent previously unannotated protein isoforms.

To validate the translation of RD-ORFs, we utilized tandem mass tag (TMT) MS data from iNeuron cells ^37^, which exhibited the highest abundance of transcript models. The peptide mass spectra from iNeuron cells were queried against the UniProt catalog (2024 April release) and two versions supplemented with polypeptide sequences encoded by RD-ORFs from DRS or CPS. After excluding peptides overlapping UniProt annotations, we identified 48 peptides exclusively matching DRS-derived RD-ORFs compared to 6 private to CPS (**Figure 2e**), representing 34 and 5 unique predicted protein sequences, respectively (**Figure 2f**). This 6.8-fold difference in recall is partially explained by the roughly 2-fold higher number of RD-ORFs discovered by DRS compared to CPS (**Figure 2g**), resulting in a 3.4-fold higher validation rate (**Figure 2h**). These findings highlight the sensitivity of DRS in combination with MS to validate transcript model translation and to potentially discover novel translated regions, or translons ^38^.

#### DRS identifies putative ORFs from medically relevant genes

Several DRS-derived RD-ORFs map to medically-relevant genes in key neuropathological gene sets: Alzheimer’s Disease (AD), Neurodegenerative Diseases (NeuDegen), and Neuropsychiatric Disorders (NeuPsy) (**Figure 2i**) ^6,39^. DRS demonstrated coverage across all three neuropathological gene sets and revealed cell type-specific distribution of RD-ORFs, which highlights its potential to characterize the molecular components of disease. Thus, we extended the use of DRS to identify medically-relevant ORFs from the post-mortem frontal cortex of eleven AD patients (**Supplemental Table 1**), which revealed a 3.3-fold overrepresentation of ORFs in neuropathology-associated genes compared to all other expressed genes (CPM > 1; two-sided Fisher exact test *p*-value = 4.03e-3). This observation may be partially explained by 1.6-fold higher expression of neuropathology-associated genes among AD brains (two-sided Mann-Whitney *U* test *p*-value = 8.93e-57). Across both iPSC-derived samples and AD brain, ORFs mapping to *APP* and *MAPT* were of particular interest given their roles in AD pathology and as therapeutic targets. Based on *BLASTp* alignment to the RefSeq protein catalog, the two predicted polypeptide sequences from *APP* ORFs matched reference protein isoforms: the isoform i precursor (733 a.a.; RefSeq subject = NP_001191231.1) in iMicroglia and the isoform k precursor (714 a.a.; RefSeq subject = NP_001372182.1) in AD brain (**Figure 2j**). These isoforms are distinct from the three canonical major isoforms APP695, APP751 and APP770 ^40^. For *MAPT*, three predicted polypeptides were detected matching reference protein isoforms: the isoform 8 (2N3R; 410 a.a.; RefSeq subject = NP_001190181.1) in AD brain, the isoform 4 (0N3R; 352 a.a.;RefSeq subject = NP_058525.1) in iNeuron, and the isoform X16 (418 a.a.; RefSeq subject = XP_047292037.1) in iAstrocyte (**Figure 2k**). The 0N3R isoform, which lacks N-terminal repeats but includes three microtubule-binding repeats, is the predominant form found in iNeurons. This finding is notable because the 0N3R isoform is also characteristic of the developing brain, which aligns with the observation that iPSC-derived neurons exhibit a fetal transcriptional profile. ^41,42^. The transcript models for *APP* and *MAPT* ORFs exhibited alternative UTRs when compared to annotated transcripts, which is important because these regions are critical regulators of protein production by controlling transcript stability, localization and translation rate ^43^. Taken together, these findings highlight the capability of DRS to identify unannotated transcript variants that encode medically-relevant ORFs, which share sequence homology with externally-validated reference protein isoforms.

#### Validation of brain-derived ORFs using ROSMAP MS data

To validate translation of brain-derived predicted ORFs, we utilized an external TMT MS dataset from the Religious Orders Study and Memory and Aging Project (ROSMAP) project ^44^. Peptide mass spectra from the post-mortem dorsolateral prefrontal cortex of eight AD patients were queried against UniProt databases supplemented with DRS-derived predicted polypeptides from our internal AD brain samples. After excluding peptides mapping to UniProt annotations, 89 unique peptides were discovered, corresponding to 49 genes. STRING database analysis (interaction score >= 0.4) revealed that these genes are associated with neuron-specific biological processes (**Figure 2l**) ^45,46^. For example, validated translons of *GRIA4* and *SYT1* form a nexus of interactions, potentially related to the role of these genes in synaptic signaling (**Figure 2m**) ^47,48^. These results highlight the potential importance of DRS-derived ORFs and their translons in neuronal processes associated with AD.

### Section 3: Calibrated DRS enables reliable RNA modification detection

#### RNA modification detection accuracy is dependent on modification type

DRS provides a unique advantage over CPS by enabling the detection of modified RNA bases. Using the RNA004 DRS chemistry with the *Dorado* v1.0.0 basecaller, we determined aggregate modification detection accuracy by contrasting the modification probability score distributions of unmodified ERCC/SIRV and SH-SY5Y genomic sites. This modification false discovery rate (modFDR) is unique to each RNA modification type and is sensitive to the probability score threshold used to classify RNA modification sites. For m6A, we observed a significant drop in modFDR at sites with modification probability scores exceeding 0.95 (**Figure 3a**), consistent with prior studies using *in vitro* transcription (IVT) ^49–51^. These high-confidence modification sites were subsequently used for all further analyses. Next, we validated DRS m6A sites with established immunoprecipitation-based ^13,14^ and chemical-conversion ^52^ methods. Using the EpiPlex immunoprecipitation protocol ^53^, we observed substantial DRS-derived m6A site coincidence, with an adapted fraction-in-peak (FRiP) score indicating 57% overlap (**Figure 3b**). In addition, base-resolved m6A stoichiometry determined by DRS showed strong correlation with results from the GLORI chemical-conversion protocol (Pearson *r* = 0.79, *p*-value = 0.00) (**Figure 3c**). Furthermore, SH-SY5Y transcript reads exhibited significantly higher m6A probabilities at the central base of DRACH motif 5-mers compared to that from unmodified ERCC/SIRV controls (**Figure 3d**) ^13,14^, confirming the ability of DRS to accurately detect biologically-derived m6A.

**Figure 3.**
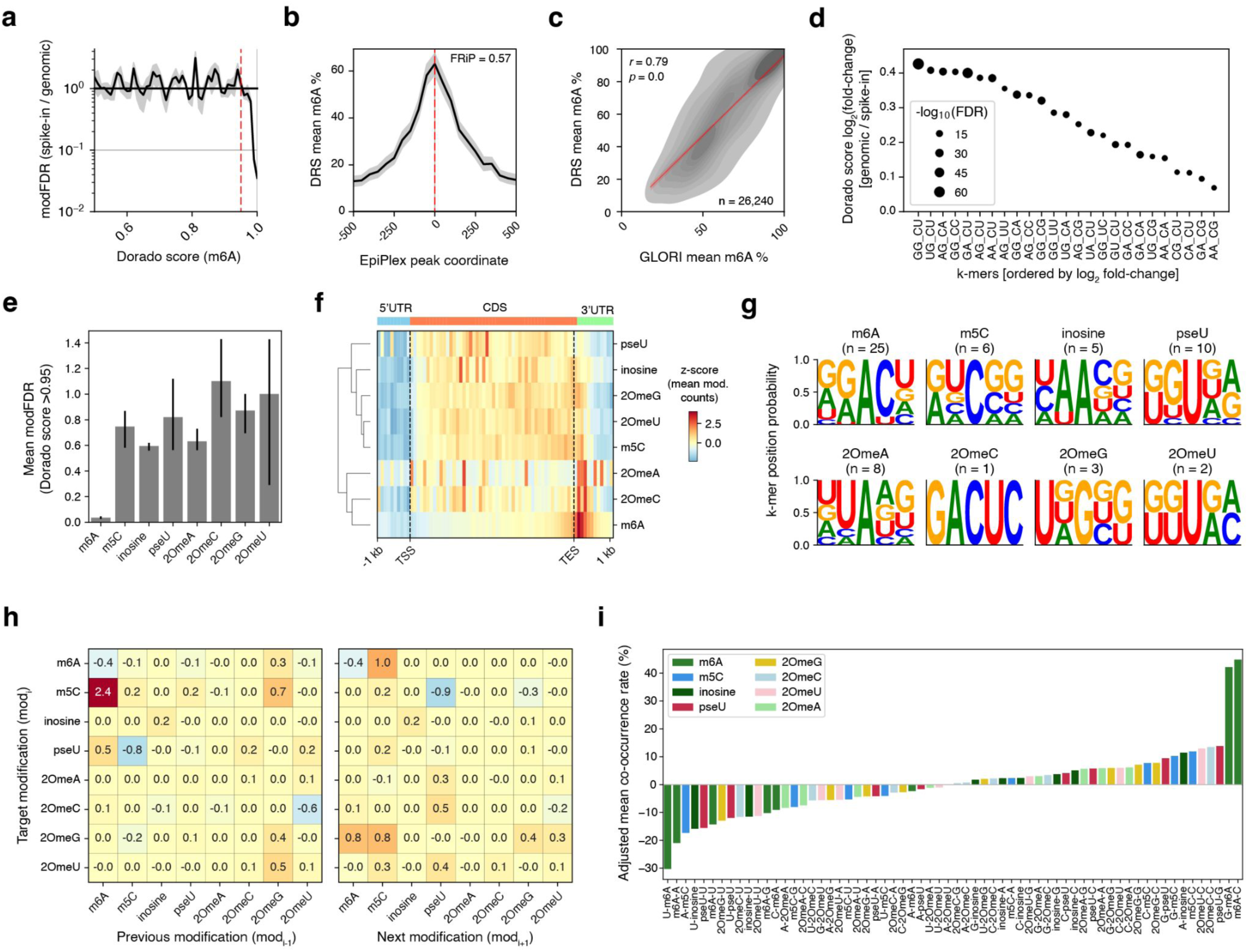
RNA modification detection performance. **a**) Line plot showing mean modification false discovery rate (modFDR) of m6A as a function *Dorado* score (*i.e.* probability of modification). The shaded area represents the 95% confidence interval, across replicates (n = 3). **b**) Line plot displaying DRS-derived m6A distribution (n = 3) relative to orthogonal m6A peak coordinates using EpiPlex (n = 1). The fraction-in-peak (FRiP) value summarizes genome-wide m6A overlap between DRS and EpiPlex. **c**) 2-dimension density plot depicting the m6A stoichiometries between DRS (n = 3) and GLORI (n = 1). Pearson correlation coefficient, associated P value and sample size are shown. **d**) Scatter plot showing 5-mers with significantly higher m6A modification probabilities, across replicates (n = 3), in the genomic context compared to an unmodified synthetic random sample. The 5-mers are ranked based on the magnitude of modification probability fold change. The central base denoted as “_” is m6A. One-sided Welch’s *t*-test FDR was used to determine significance. Point sizes are scaled to the negative log_10_ transform of FDR. **e**) Bar plot displaying the mean modFDR for modification sites, across replicates (n = 3), with a Dorado score of greater than 0.95, with error bars representing the 95% confidence intervals. **f**) Heatmap illustrating the mean metagene distribution of modifications, across replicates (n = 3). Metagene bins are color-coded by the *z*-score of mean modification percentage. The modifications are grouped based on hierarchical clustering. **g**) Sequence logos of composite 5-mers with a central modification, based on the position frequency matrices of 5-mers exhibiting significantly higher genomic modification probabilities. The respective number of significant 5-mers is shown. **h**) (left) Heatmap showing the adjusted mean co-occurrence rate (%) for the target modified base relative to an adjacent previous modification, across replicates (n = 3). (right) Similar to before, except relative to an adjacent next modification. **i**) Barplot displaying adjusted mean co-occurrence rate (%) for dinucleotides that contain a mix of modified and unmodified bases. Bars are color-coded based on the modified base identity within the dinucleotide.

Beyond m6A, the *Dorado* basecaller can characterize seven additional RNA modifications: m5C, pseU, inosine, and 2’-O-methylation of canonical RNA bases (2OmeA, 2OmeC, 2OmeG, 2OmeU). However, these modifications exhibited substantially higher modFDRs compared to m6A (**Figure 3e**), consistent with prior literature ^54^. Despite this, we proceeded to evaluate aggregate modification site distribution across transcript regions, which revealed two high-level groupings: m6A, 2OmeA, and 2OmeC localize to the transcription end site (TES) and 3’-untranslated region (3UTR) while the remaining modifications are more evenly distributed along the protein-coding region (CDS) (**Figure 3f**). Next, we identified the local sequence motif of each RNA modification by aggregating significantly modified 5-mers into sequence logos (one-sided Welch’ *t*-test FDR < 0.05; **Figure 3g and EDF 3**). Gratifyingly, several RNA modification sequence logos were consistent with prior reports ^55–57^, such as the DRACH motif at m6A sites. In addition, we observed a G-rich motif at m5C sites, which aligns with the sequence preference of a m5C writer enzyme, NSUN2 ^58^. Taken together, DRS modification calls exhibited biologically relevant patterns in aggregate analyses; while, single-molecule accuracy was less reliable and modification-specific.

#### DRS facilitates adjacent RNA modification detection

We set out to comprehensively characterize adjacent RNA base modifications across those detected by *Dorado*. In triplicate, we computed the mean co-occurrence rate for eight modifications resulting in 64 couplings while considering both orientations (*i.e.* 5’-mod_x_-mod_y_-3’ versus 5’-mod_y_-mod_x_-3’), equal to 128 unique adjacent modification pairs. To correct for sequence bias inherent to the modification calling model, the co-occurrence rates were baseline-normalized using unmodified ERCC/SIRV reads, resulting in an adjusted rate. Basecaller cross-reactivity is another source of errors, which we anticipate can be mitigated with improved sampling of diverse sequence contexts containing adjacent modification pairs during model training ^11^. Acknowledging this potential limitation, our analysis revealed a substantial positive co-occurrence rate of 2.4% when m5C is preceded by m6A (**Figure 3h**), consistent with previous reports ^11,12^. Interestingly, this relationship is asymmetric, as m5C followed by m6A showed no association. A case of adjacent modification antagonism was detected for m5C followed by pseU, which showed a negative co-occurrence rate. This relationship is also asymmetric, as the reverse sequence does not show this antagonistic effect. While the mean co-occurrence rates for adjacent modifications are modest when compared to those of mixed dinucleotides (composed of modified and unmodified bases; **Figure 3i**), they could still hold biological significance at specific loci. These observations provide initial insights into the complex interactions of RNA modifications.

### Section 4: The interplay between m6A and the ribosome

#### Integration of ribosome positions with m6A sites reveals epitranscriptomic interaction with protein translation machinery

To investigate how m6A epitranscriptomic signals influence protein translation in a neurobiological context, we analyzed iNeurons using multiple analytical approaches (**Figure 4a**). We examined how m6A marks intersect with ribosome positions, aiming to elucidate epitranscriptomic mechanisms bridging transcription and translation. Recent studies have suggested that m6A sites can modulate tRNA-mRNA interactions within the ribosome A-site, leading to mRNA decay ^59–61^. Extending these findings to iNeurons, we found a modest but significant negative correlation between m6A levels in the CDS and transcript abundance (Pearson *r* =-0.21, FDR = 0.00; **Figure 4b**). At single-nucleotide resolution, elevated m6A levels were observed within predicted A-sites determined by Ribo-seq (**Figure 4c**). Further analysis revealed preferential position-specific m6A modification of codons mapped to ribosome A-sites, in some cases diverging from codon usage predicted from the DRACH motif ^14,55^ (**Figure 4d,e,f**). For A-site codons with m6A modification at the first nucleotide position (A_n1), we observed excellent correlation with predicted codon usage (Pearson *r* = 0.98; **Figure 4g**). In contrast, m6A modification of the second and third positions (A_1_ and A_2_) was associated with codon frequencies that deviate from predicted proportions (Pearson *r* = 0.86 and 0.60, respectively; **Figure 4h,i**). This m6A-modified A-site codon profile leads to an altered frequency of amino acid usage, showing a preference for aspartate (D), glycine (G), and arginine (R) (**Figure 4j,k,l**).

**Figure 4.**
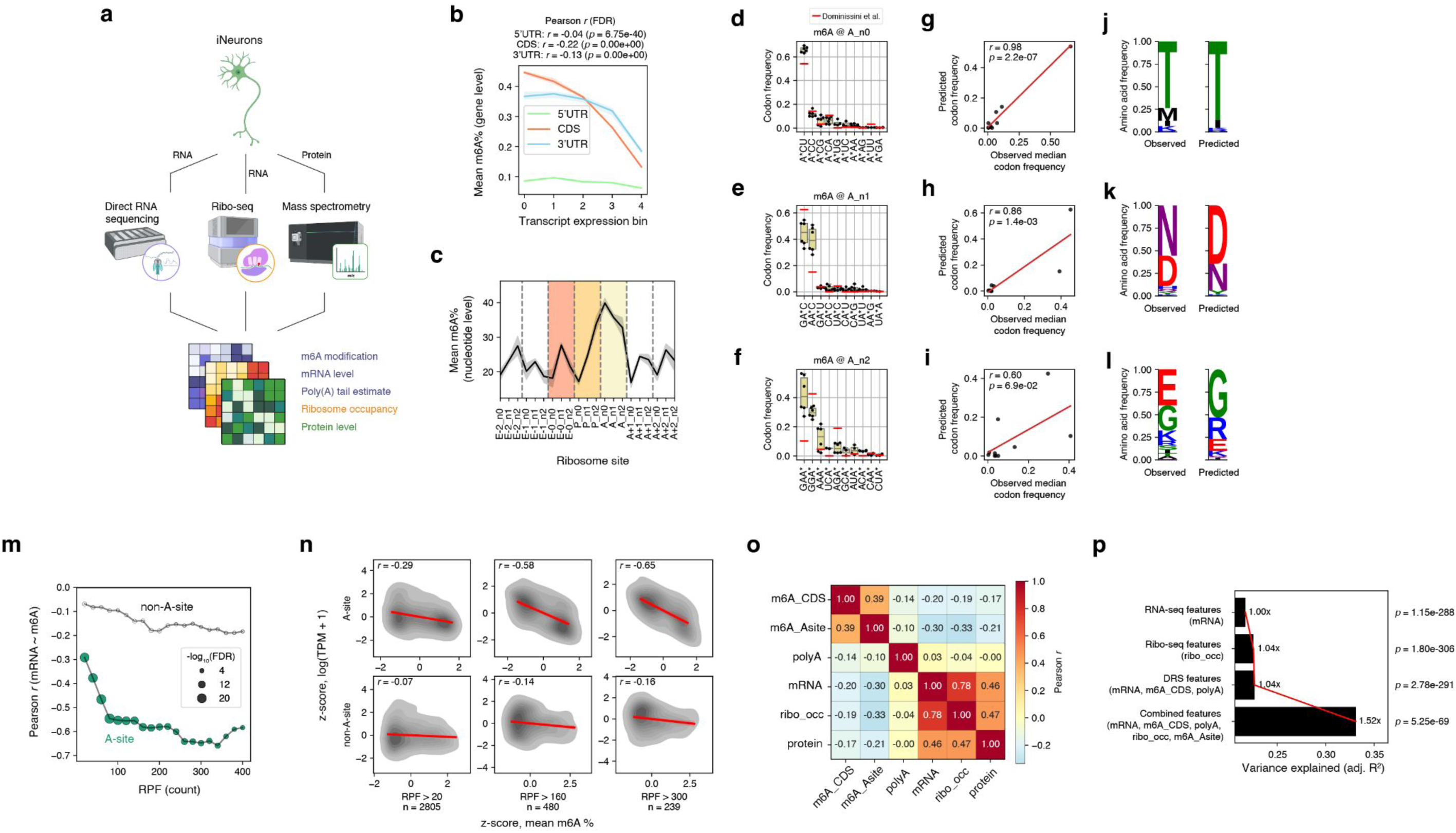
Integrative multi-omic characterization of m6A in iNeurons. **a**) Schematic representation of the integrative multi-omic characterization of m6A in iNeurons. **b**) Line plot showing the *z*-score of mean m6A percentage versus relative transcript abundance expression bins, across replicates (n = 3), disaggregated by m6A location in either UTR5, CDS or UTR3. The shaded area represents the 95% confidence interval. Pearson correlation coefficient and associated FDR are shown. **c**) Line plot depicting the distribution of m6A (DRS: n = 3), along ribosome-bound mRNA fragments (Ribo-seq: n = 2). The ribosome site on the x-axis refers to the three major ribosomal sites: Exit site (E), Peptidyl site (P) and Aminoacyl site (A). The shaded area represents the 95% confidence interval. **d**) Boxplots displaying the codon usage proportion within the ribosomal A-site, where m6A is detected at the first nucleotide position. Each point represents a unique comparison between DRS and Ribo-seq replicates (DRS: n = 3, Ribo-seq: n = 2), with proportions based on the colocalization of m6A modification sites and ribosome-protected fragments. The codons are ranked based on median usage proportion. The asterisk indicates the site of m6A modification within the codon. Predicted codon usage based on DRACH motif shown as red line from Dominissini *et al*.^14^ **e**) Similar to before, except for the second nucleotide position of an A-site codon. **f**) Similar to before, except for the third nucleotide position of an A-site codon. **g-i**) Linear regression plot of observed median codon frequency versus predicted codon frequency. Pearson correlation coefficient and P value are shown. **j-l**) Sequence logo of the corresponding amino acid usage proportion derived from both observed and predicted codon usage. **m**) Scatter plot illustrating the Pearson correlation between mRNA abundance and m6A modification level (DRS: n = 3, Ribo-seq: n = 2), as a function of ribosome occupancy, measured by ribosome-protected fragment (RPF) read depth. Points colored yellow indicate m6A detected in A-sites. The size of points scales with the negative log_10_ transform of the FDR associated with each Pearson correlation. **n**) Kernel density estimation plots with regression line in red showing gene-level standardized (*z*-score) mean m6A % versus scaled expression. Transcripts with predicted ribosome A-sites overlapping m6A sites are summarized in the top sub-panel series labeled “A-site”; whereas, those transcripts with mutually-exclusive predicted ribosome A-sites and m6A sites are summarized in the bottom sub-panel series labeled “non-A-site”. RPF depth threshold, number of genes and Pearson r are shown. **o**) Heatmap showing all pairwise Pearson correlations between measurement types. For measurements with replicates (DRS: n = 3, Ribo-seq: n = 2, TMT MS: n = 1), mean values were used and preprocessing standardization to zero mean and unit variance was performed before correlation. The number of genes in each comparison is dependent upon the overlap of detected genes between measurement types in a given pair. **p**) Boxplot displaying adjusted coefficient of determination (R^2^), or variance explained, for a series of multivariate linear regression models for combinations of outputs from DRS, Ribo-seq, and MS. Fold-change calculations relative to “RNA-seq features” are shown at the end of each bar. Global *F*-test P values are shown along the right y-axis.

#### DRS sheds light on m6A-associated mRNA decay

Next, we show that m6A levels are negatively correlated with transcript abundance across a range of ribosome occupancy levels, especially when m6A is in an A-site, which leads to a 3-to 6-fold increase in the correlation coefficient magnitude (**Figure 4m,n**). This difference was most pronounced at loci with approximately 100 ribosome-protected fragment counts, indicating substantial ribosome occupancy. These findings are consistent with the CDS-m6A decay (CMD) model, in which m6A in A-site codons promotes mRNA decay, potentially *via* recruitment of the CCR4-NOT complex that facilitates poly(A) tail deadenylation ^59,61^. To investigate this in the context of iNeuron cells, we employed DRS to estimate mean poly(A) tail lengths per transcript and analyzed their relationship with both m6A levels and mRNA abundance. Consistent with CCR4-NOT-mediated deadenylation of m6A-modified transcripts, we observed a slight negative correlation between poly(A) tail lengths and m6A deposition within the transcript CDS and at A-site codons (Pearson *r* =-0.14 and-0.10, respectively; **Figure 4o**). Surprisingly, we find a very weak dependence between poly(A) length and mRNA abundance (Pearson *r* = 0.03), a finding that has been corroborated in an analogous neuronal cell differentiation model ^62^. Given the potential link between poly(A) tail length and translational efficiency ^63^, we compared our findings with reported protein quantification levels from iNeurons ^37^, which revealed no correlation between poly(A) tail lengths and protein abundance. These findings suggest that additional mechanisms beyond m6A-templated deadenylation of transcripts are responsible for the repression of both mRNA and protein expression.

#### Ribo-seq and DRS feature integration improves protein expression prediction

We next employed multivariate linear regression to assess the contribution of DRS and Ribo-seq features in predicting protein expression. First, we established the baseline association between mRNA abundance and protein expression, which is the implicit basis of typical RNA-seq applications, revealing a modest adjusted *R*² of 0.217 (**Figure 4p**). Ribosome occupancy determined by Ribo-seq showed only a slightly higher adjusted *R*² of 0.225. The adjusted *R*² for DRS, which simultaneously provides mRNA abundance, CDS-specific m6A percentage, and poly(A) tail length, was 0.227. Both Ribo-seq and DRS explained a similar proportion of the protein expression variance, with each technique improving upon mRNA abundance alone by a factor of 1.04. Notably, integration of Ribo-seq and DRS, thus enabling the measurement of m6A levels within ribosome A-sites, substantially improved explanatory power by 1.52-fold compared to baseline. The observed synergy of combining transcriptomic, epitranscriptomic, and translatomic data for improved protein abundance predictions demonstrates the value of this integrative approach in advancing our understanding of protein expression regulation.

### Section 5: m6A profiles are distinct across cell types and disease stages

#### DRS generates cell type-specific transcriptome-wide m6A profiles

We next sought to understand the distribution of m6A within the CDS of transcripts throughout the transcriptome of iPSC-derived cell types in addition to eleven AD brain samples. Measuring the mean m6A percentage within CDS regions (%m6A*_CDS_*) across all genes for each sample revealed substantially elevated m6A deposition in iN (two-sided Mann-Whitney *U* test *p*-value = 1.13e-3; **Figure 5a**). Next, we perform principal component analysis (PCA) using gene-level %m6A*_CDS_*, which revealed segregation between AD samples and iPSC-derived brain cell types (**Figure 5b**). Furthermore, we observed the expected partitioning among neurons and glial cells. Gene set enrichment analysis (GSEA) was performed using the feature loadings of the first two principal components, which yielded expected cell type-specific biological processes based on differential m6A levels (**Figure 5c**).

**Figure 5.**
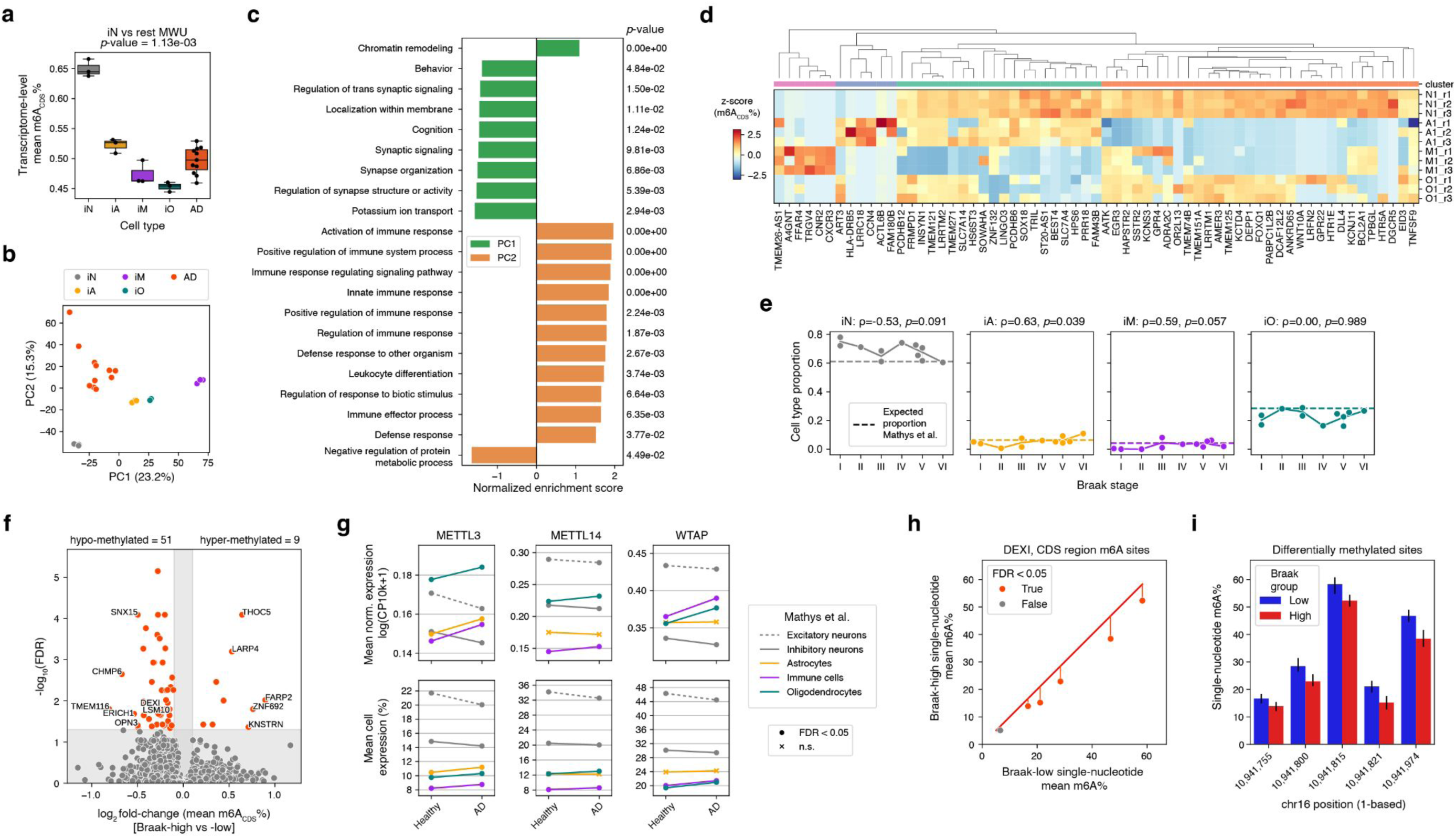
Cell type-specific and disease stage-specific m6A profiles. **a)** Boxplot depicting transcriptome-level mean percent m6A in the CDS (%m6A_CDS_) across iPSC-derived cell types and AD samples, color-coded by sample type. Two-sided Mann-Whitney *U* test P value shown for iN (n = 3) vs rest (n = 20). **b)** Scatterplot showing PCA embeddings using highly variable gene-level %m6A_CDS_, color-coded by sample type in legend. **c)** Barplot displaying significantly enriched Gene Ontology Biological Process terms based on PC loadings. **d)** Heatmap depicting normalized %m6A_CDS_ for selected genes versus iPSC-derived cell type sample. Genes are grouped by hierarchical clustering with associated color-coded “cluster” annotation below dendrogram. **e)** Scatterplot showing SVR estimates of cell type proportion separated by Braak stage, separated in subpanels based on cell type. The mean proportion is shown along the line connecting each Braak stage. A horizontal dashed line displays the expected cell type proportion based on Mathys *et al.* ^65^. Color corresponding to cell type. Spearman ⍴ and P values are shown. **f)** Volcano plot displaying differentially methylated genes identified using a binomial GLM based on %m6A_CDS_, contrasting Braak-high (stages IV-VI; n = 6) and Braak-low (stages I-III; n = 5). Genes colored orange exhibit an absolute log_2_ fold change of > 0.01 and FDR < 0.05. The total number of hypo-and hyper-methylated genes is shown. **g)** Line plots depicting m6A writer complex component expression between health and disease, disaggregated by major cell type annotation, based on data from Mathys *et al.* ^65^. The top subpanel series shows mean normalized expression: natural log of counts per 10,000 (CP10k) + 1. The bottom subpanel series shows the mean percent of cell expressing. Major cell type annotations are color-coded and neuron subtypes are distinguished by solid versus dashed lines. Markers at line termini reflect statistical significance: Mann-Whitney *U* test FDR < 0.05 or not significant (*n.s.*). **h)** Scatterplot showing single-nucleotide mean % m6A, comparing Braak-low (stages I-III; n = 5) versus Braak-high (stages IV-VI, n = 6) samples. Binomial GLM statistical significance ( FDR < 0.05 versus *n.s.*) is shown in color-coding. The diagonal line depicts equality between Braak groups. Vertical lines originating from the diagonal to each significant point highlights deviation from equality. **i**) Barplot displaying mean single-nucleotide %m6A at each significant differentially-modified position, color-coded by Braak group. Error bars represent data range.

#### Identification of physiologically-relevant differentially methylated genes

With the expectation that iPSC-derived cells may exhibit some non-physiological m6A deposition, we sought to identify a parsimonious subset of genes with %m6A*_CDS_* levels that recapitulate bulk AD brain. Bulk deconvolution methods, initially developed for RNA-seq data, were adapted for m6A analysis. The goal was to identify physiologically-relevant gene subsets whose methylation profiles are consistent across both iPSC-derived cells and AD brain samples. Toward this end, we modified the CIBERSORT deconvolution approach based on support vector regression (SVR)^64^ to ingest cell type-specific m6A signatures and estimate cell type proportions of bulk AD brain samples. Candidate gene subsets were assessed by comparing SVR estimates with major cell type proportions published in a snRNA-seq prefrontal cortex atlas containing 427 ROSMAP study participants ^65^. This study reported approximately 58.2% neurons, 6.3% astrocytes, 3.6% immune cells (predominately composed of microglia), 27.3% oligodendrocytes, and the remaining 4.6% of cells were oligodendrocyte precursor and vascular cells. To efficiently explore gene subsets that recapitulate bulk AD brain, we implemented a genetic algorithm that iteratively selects for genes that minimize SVR error. From 250 random initializations of this genetic algorithm, we identified 63 significantly over-represented genes clustered into four groups reflective of iPSC-derived brain cell types (one-sided exact binomial test FDR < 0.05; **Figure 5d**) that together yielded cell type proportion estimates roughly aligned with expectation (**Figure 5e**). Interestingly, we observed a trend between predicted cell type proportions and Braak stage, despite limited statistical power due to small sample size. This trend involved a decrease in the proportion of induced iN (Spearman ⍴ =-0.53, *p*-value = 0.091) and an increase in the proportions of iA (Spearman ⍴ = 0.63, *p*-value = 0.039) and iM (Spearman ⍴ = 0.59, *p*-value = 0.057). This shift in cell type composition is consistent with that observed in reactive gliosis among symptomatic AD patients ^66^. Overall, this workflow demonstrates the utility of bulk deconvolution with cell type-specific %m6A*_CDS_* signatures for identifying physiologically-relevant and differentially-modified genes, which recapitulate the m6A profile observed in AD brain.

#### Differentially methylated genes can distinguish between early and late AD

Next, we sought to focus on the m6A modification in the CDS of AD samples, specifically looking at early compared to late stage disease. Therefore, we performed differential methylation analysis for Braak-low (stages I-III) versus Braak-high (stages IV-VI) samples using a binomial generalized linear model (GLM) with deviance-based dispersion adjustment, akin to the statistical framework developed for DNA methylation analysis ^67–69^. This analysis identified 9 hyper methylated genes and 51 hypomethylated genes (GLM FDR < 0.05; **Figure 5f**). This observation is consistent with reports of general hypomethylation in AD brains compared to healthy and early AD brains ^21,70^. Furthermore, reported downregulation of the m6A writer complex components METTL3, METTL14 and WTAP during AD progression led us to determine if such dysregulation is evident in our 11 AD samples ^71^. Differential gene expression analysis contrasting Braak-low versus Braak-high samples revealed no significant change in the m6A writer complex component expression, which we reasoned might be due to the limited statistical power afforded by our small sample size. Therefore, we queried the aforementioned ROSMAP snRNA-seq dataset composed of 2.3 million cells ^65^. By comparing healthy brains versus those with a pathologic diagnosis of AD, we observed cell type-specific downregulation of METTL3, METTL14 and WTAP in neuronal subtypes concomitant with upregulation in immune cells and oligodendrocytes (Mann Whitney *U* test FDR < 0.05; **Figure 5g**). While indirect, these findings suggest that neuron-specific downregulation of m6A writer machinery might be responsible for the observed global hypomethylation in our 11 AD samples.

#### DRS enables single-nucleotide differential methylation analysis across AD groups

Of the genes found to be hypomethylated in Braak-high samples, *DEXI* showed the largest absolute change, with its mean %m6A*_CDS_* reduced by 1.05%. (GLM FDR < 0.05; **EDF 4**). To determine differential m6A levels at single-nucleotide resolution, we redeployed the binomial GLM method adapted for individual modification sites, which revealed several significant hypomethylated positions within *DEXI* (FDR < 0.05; **Figure 5h**). Specifically, the modification site chr16:10,941,974 corresponding to the second nucleotide of an aspartate codon (GAC), showed the greatest reduction in mean %m6A, lower by an average of 8.3% among Braak-high samples (**Figure 5i**). Interestingly, all five hypomethylated sites identified within the CDS of *DEXI* are located in aspartate codons. As we discovered in iNeurons, m6A-modified aspartate codons are overrepresented within the ribosomal A-site, which contrasts with predicted codon usage based on the canonical DRACH motif. These analyses emphasize the importance of using DRS for single-nucleotide modification detection and quantification, which enables the exploration of how m6A position, level, and multiplicity within a transcript impacts function.

## Discussion

DRS has emerged as a powerful tool for transcriptomic and epitranscriptomic studies, offering the unique ability to sequence native RNA and detect modifications with single-nucleotide resolution. In this study, we utilized DRS to explore the molecular underpinnings of AD and related *in vitro* models, demonstrating its superior accuracy and sensitivity compared to CPS. This heightened sensitivity suggests that DRS provides a more comprehensive and unbiased view of the transcriptome, circumventing biases inherent in cDNA-based methods. DRS also proved instrumental in identifying transcript models with protein-coding potential, some of which were shown to be translated using MS-based detection.

Using the *Dorado* basecaller coupled with a false-discovery minimization strategy calibrated with unmodified controls, we accurately identified m6A modifications at single-nucleotide resolution. Importantly, DRS-derived results recapitulate the canonical DRACH motif of m6A and were corroborated using established immunoprecipitation-based and chemical-conversion methods. Beyond m6A, aggregate analyses revealed biologically meaningful patterns for other RNA modifications, despite lower single-molecule accuracy. Notably, DRS uncovered interactions between adjacent modifications, such as the elevated co-occurance of m5C preceded by m6A, and antagonism between m5C and pseudouridine. The ability to detect multiple RNA modifications simultaneously highlights the potential of DRS to decipher the “epitranscriptomic code” that regulates RNA function.

DRS also provided key insights into the relationship between m6A and protein translation in iNeurons. Using an integrative multi-omic approach, combining DRS, Ribo-seq and MS, we observed that both transcript and protein abundances were negatively correlated with m6A levels, particularly when the modification is located at a ribosome A-site. These results are consistent with recent research linking m6A-templated mRNA decay and translational repression. The ability of DRS to also estimate the poly(A) tail length of transcripts enabled a deeper mechanistic analysis into the role of transcript polyadenylation in this epitranscriptomic regulatory model. While m6A deposition was modestly associated with shorter poly(A) tail lengths, we observed no relationship between poly(A) tail length and mRNA or protein abundance, suggesting that m6A-mediated repression involves additional mechanisms beyond tuning poly(A) tail lengths. Rather, previous reports implicate sequestration of translationally repressed mRNAs to P-bodies, where transcript degradation occurs *via* 7-methylguanosine decapping followed by XRN1-mediated 5’-to-3’ exonucleolytic decay ^59,72^.

Furthermore, our DRS-based analysis of m6A in iPSC-derived cell types and AD brain samples revealed cell type-specific m6A profiles, which allowed for the deconvolution of bulk AD brain samples. This approach identified a subset of 63 genes whose %m6A*_CDS_* levels recapitulated expected cell type proportions in bulk AD brain. Intriguingly, the estimated cell type composition trended with Braak stage, indicating a shift toward reactive gliosis in symptomatic AD. Moreover, differential methylation analysis comparing early (Braak I-III) and late-stage (Braak IV-VI) AD brains identified 9 hypermethylated and 51 hypomethylated genes, consistent with general hypomethylation reported in advanced AD. The greatest reduction in %m6A*_CDS_* was observed for *DEXI*, and single-nucleotide resolution analysis further pinpointed specific hypomethylated m6A sites within its CDS, all located in aspartate codons. These findings underscore the utility of DRS for detailed m6A quantification, enabling the identification of physiologically-relevant, differentially modified genes and specific m6A positions that may impact transcript function in AD.

Despite its significant advantages, this study also highlights limitations of DRS, including lower read throughput and reduced raw read accuracy compared to CPS. And, recent systematic benchmarking indicates superior performance of CPS over DRS on certain metrics, dependent on sample type ^5^. In our study, one area of relative advantage for DRS is in longer read length distributions, which facilitates enhanced transcript sensitivity. Recent work to improve the detection of long isoforms using DRS shows promise in further enhancing transcript sensitivity ^73^. However, we expect that continued advancements in cDNA library preparation will increase read lengths and reduce PCR bias (*e.g.* standardized use of unique molecular identifiers to deduplicate reads), thus narrowing the performance gap between DRS and CPS. For detecting RNA modifications, we acknowledge that single-molecule accuracy was highly variable across modification types. We anticipate that advancements in basecalling models will both enhance accuracy and broaden the range of detectable modifications. Indeed, a recently released deep learning framework for modification detection called DeepRM claims near-perfect accuracy at single-molecule resolution ^74^. Furthermore, DRS enables the detection of co-occurring RNA modifications, but basecalling cross-reactivity errors remains a limitation ^11,12^. Therefore, appropriate basecalling model design and training are necessary to mitigate the risk of false positives. And, while bulk brain tissue analysis with DRS has yielded important insights into disease progression, a critical gap remains: the lack of scalable, sensitive and cost-effective methods applying DRS in single cells. Such methods would be useful for distinguishing between cell state expression changes versus alterations in cell proportions. For instance, observed hypomethylation in late-stage AD may be due to a smaller population of neurons, rather than an altered m6A profile of surviving neurons.

In summary, DRS is a powerful platform for transcriptomic and epitranscriptomic research, enabling the characterization of transcript isoforms and detection of RNA modifications. We demonstrated the sensitivity of DRS, in combination with MS, to discover novel translons. DRS was also used to characterize the role of m6A in regulating mRNA abundance through engagement with ribosomes, and thus enabling more accurate prediction of protein translation. As the technology advances, DRS is poised to uncover new layers of RNA biology as well as provide a deeper understanding of the molecular components underpinning neurodegenerative diseases and other complex disorders.

## Materials and Methods

### iNeurons

NGN2 iPSCs (iP11N cells batch 7884 from CellCentral) were differentiated into neurons using the protocol as described in Shan *et al* ^37^. Briefly, iPSCs were induced with DMEM/F12 (11320033), supplemented with B27 without VA (Thermo Fisher 12587010), N2 Max (R&D Systems, AR009), Glutamax (Thermo Fisher 35050061), NEAA (Thermo Fisher 11140050), SB431542 (10 uM, Cell Signaling 14775S), XAV939(2 uM, StemCell Technologies 72672), Noggin (100 ng/mL, Mitenyl 130-103-456), DAPT (10 uL) and Doxycycline (3 ug/mL,Takara Bio 631311). Following induction and selection with Ara C (5 uM, Sigma C1768) for 3 days, cells were replated into 10cm plates at 6M cells per plate on Day 8. Cells were then maintained in Naurobasal Plus Medium (Thermo Fisher A35829-01), supplemented with B27 without VA (Thermo Fisher 12587010), Glutamax (Thermo Fisher 35050061), BDNF (20 ng/mL, R&D Systems 248-BDB-010), GDNF (20 ng/mL, R&D Systems 212-GD/CF), Ascorbic Acid (200 uM, StemCell Technologies 72132), cAMP (500 uM, Millipore Sigma D0627), DAPT (10 uM, until Day 14), Dox (2 ug/mL until Day 14) from Day 8 to Day 28 for maturation. The cells were dissociated on Day 28 with 3 units/mL papain (LK003150) at 37C for 10 min. Cells were gently lifted and spun down at 300g for 5 min.

### iMicroglia

Hematopoietic progenitors generated as described by Guttikonda *et al.* ^75^ and Zhang *et al.* (under review) were expanded between days 10–14 through incremental media supplementation under defined cytokine conditions to support myeloid differentiation. On day 14, suspension cells were collected and transferred to IL-34– and M-CSF–containing media, then plated onto fibronectin-coated vessels. Between days 16–18, cultures were supplemented to maintain loosely adherent and suspended populations. On day 20, adherent and non-adherent cells were collected and pooled, then either re-plated in differentiation media containing IL-34, M-CSF, and TGF-β1 or cryopreserved. From day 22 onward, cells were transitioned through reduced-serum conditions into fully defined serum-free media with staged increases in TGF-β1 to promote microglial maturation. Media changes were performed every other day, and cultures were maintained in IL-34, M-CSF, and TGF-β1 until maturation. Mature iMicroglia collected at day 31 were used for downstream profiling in this study.

### iAstrocytes

Astrocyte precursor cells (APCs) were generated based on previously described methods {Russo, 2018 #378}. Briefly, iPSCs were differentiated into neural progenitor cells (NPCs) using the STEMdiff SMADi Neural Induction Kit (StemCell Technologies; Cat. No. 08581) according to the manufacturer’s instructions. Neural rosettes were manually isolated and expanded in NPC medium (DMEM/F12, Neurobasal, GlutaMAX, B27, N2) supplemented with FGF2, EGF, and BDNF until confluent. NPCs were then dissociated into single cells and seeded into AggreWell 800 plates (StemCell Technologies; Cat. No. 34815) at a density of 10,000 cells per microwell to form spheroids. Spheroids were cultured in NPC medium for one week, followed by a gradual transition over two weeks to astrocyte growth medium (Lonza; Cat. No. CC-3186) using partial media replacement. After three weeks, spheroids were transferred to Geltrex-coated plates at a 1:6 split ratio and maintained for an additional four weeks in an astrocyte growth medium without re-plating. By the end of this period, cells exhibited astrocyte precursor characteristics, as indicated by expression of lineage markers including GFAP, S100B, and AQP4. APCs were expanded for up to two additional passages, dissociated into single cells, and cryopreserved. Cells were fed three times per week at all stages except prior to the NPC stage. For further maturation, APCs were cultured for an additional two weeks in ScienCell Astrocyte Maturation Medium without FBS (ScienCell; Cat. NO. 1801) and then collected for downstream analyses.

### iOligodendrocytes

Induced pluripotent stem cells (Cell line PGP1; Coriell cell line GM23338) were differentiated into the oligodendrocyte lineage cells using a modified 20-day directed differentiation protocol, adapted from established methods ^76^ ^77^. Briefly, iPSCs were initially maintained on iMatrix-511 (Takara, Cat. Num. T303) coated plates. Neural induction was initiated on matrigel coated plates on Day 0 (DIV 0) by switching the basal medium to N2B27 supplemented with the Dual SMAD inhibitors SB431542 (10 µM) and LDN193189 (1 µM), and Retinoic Acid (RA, 0.1 µM). This induction cocktail was maintained until DIV 5. For Olig2+ neural progenitor cells specification (DIV 5–8), the medium was further supplemented with the Smoothened agonist SAG (10 µM). On DIV 8, cells were dissociated with Accutase and re-seeded onto Poly-L-Ornithine/Laminin-coated plates at a density of approximately 50,000 cells/cm^2^ in N2B27 supplemented with RA, SAG and 1X Revitacell. To accelerate terminal maturation, cells were transduced on DIV 9 with the doxycycline (Dox)-inducible lentiviral vector which co-expresses the Oligodendrocyte transcription factors Sox10, Olig2, and Nkx6.2 with mNeonGreen as the reporter and blasticidin for the selection. From DIV 10 onward, cells were maintained in an OL differentiation medium (OLDM) containing PDGFaa, IGF-1, HGF, NT3, T3, Biotin, and cAMP, and expression was induced by the addition of doxycycline (1 µg/mL). OPCs and immature oligodendrocytes were fed every other day with OLDM containing 1 µg/mL doxycycline and selection was started using 10 µg/mL Blasticidin from DIV16 to DIV20. Cells were harvested for sequencing or other downstream assays on DIV20.

### AD samples

Flash-frozen human brain tissue specimens were obtained from the Arizona Study of Aging and Neurodegenerative Disorders and Brain and Body Donation Program at Banner Sun Health Research Institute ^78^ under an established contract (**Supplemental Table 1**). Subsequent processing of the samples, including RNA extraction and quality control, was conducted as a service by the third-party vendor Q2 Solutions employing their standard methodologies.

### SH-SY5Y cell lines

SH-SY5Y cells were grown in RPMI-1640, 10% FBS with 2mM L-glutamine medium in a humidified atmosphere with 5% CO2 conditions. Cells were passaged at ∼ 80% confluency. For passaging, cells were pre-warmed in 1X Trypsin-EDTA in DPBS solution and incubated at 37°C for 2 minutes. Trypsin was neutralized with an equal amount of cell culture media and cells were spun down and counted using the Vi-Cell XR (Beckman Coulter). Cells were split between 1:3 to 1:10 and cultured in 100mm x 20mm Corning tissue-culture treated polystyrene dishes (Millipore, CLS430167).

### RNA Extraction

Approximately 0.35✕10^6^ - 2✕10^6^ cells were lysed using QIAzol lysis reagent and RNA was purified using the RNeasy Plus Universal Mini kit (Qiagen, 73404). RNA concentration was measured with Qubit RNA Broad Range Assay (Thermofisher Scientific, Q10210). RNA integrity was assessed using RNA ScreenTape analysis (Agilent, 5067-5576) on the 4200 TapeStation system (Agilent, G2991BA). Samples which exhibited a significant fraction of small RNA species <250 nt were subjected to an additional cleanup and size selection step using 0.6-0.7✕ SPRI with RNAcleanXP beads (Beckman Coulter, A63987).

### Long-read cDNA-PCR sequencing

500ng of total RNA was prepared using the cDNA-PCR sequencing V14 kit (Oxford Nanopore Technologies, SQK-PCS114.24, Protocol: PCB_9201_v114_revA_06Dec2023). For the SH-SY5Y cell line, total RNA was supplemented with synthetic ERCC and SIRV transcripts (Mixes E0, E1 and E2 across three independent samples) targeting approximately 1% (0.5% each) of the estimated mRNA abundance. For all samples, 14 cycles of PCR amplification was performed. Libraries were sequenced using PromethION flow cells (FLO-PRO114M) on a PromethION 24 instrument. 3 samples were multiplexed per flowcell.

### Long-read direct RNA sequencing

1μg of total RNA was prepared using the Direct RNA sequencing kit (Oxford Nanopore Technologies, SQK-RNA004, Protocol: DRS_9195_v4_revC_20Sep2023). For the SH-SY5Y cell line, total RNA was supplemented with synthetic ERCC and SIRV transcripts (Mixes E0, E1 and E2 across three independent samples) targeting approximately 1% (0.5% each) of the estimated mRNA abundance. Libraries were sequenced using PromethION flow cells (FLO-PRO004RA) on a PromethION 24 instrument.

### GLORI 2.0

Total RNA was extracted from SH-SY5Y cells using QIAzol lysis reagent and RNA was purified using the RNeasy Plus Universal Mini kit (Qiagen, 73404).. Polyadenylated mRNA was isolated using Dynabeads Oligo(dT)_25_ (Invitrogen, 61005). Approximately 200ng of PolyA+ RNA was used for input into the GLORI2.0 protocol <DOI:10.1038/s41592-025-02680-9>. Library preparation was performed using the SMART-Seq Total RNA Pico Input (w/ ZapR Mammalian) (Takara, 634357). Libraries were PEx150 sequenced on NovaSeqX+ (Illumina). GLORI-tools (https://github.com/liucongcas/GLORI-tools) (run_GLORI2.py) was used for analysis with the default settings.

### Ribosome Profiling (Ribo-seq)

Ribosome profiling was performed as previously described ^79,80^. Briefly, 10 million day 28 NGN2 iPSCs were detached using Papain (≥10 U/mg, Sigma-Aldrich) at 37°C for 10 minutes and lysed in polysome lysis buffer (20 mM Tris-HCl pH 7.4, 150 mM NaCl, 5 mM MgCl₂, 1 mM DTT, 1% Triton X-100, 25 U/mL Turbo DNase, 0.1 mg/mL cycloheximide). Lysate corresponding to 30 µg RNA was digested with 15 U RNase I (LGC Biosearch Technologies) for 45 minutes at 24°C with shaking at 400 rpm. Monosomes were isolated using MicroSpin S-400 HR columns (Cytiva) per the manufacturer’s instructions, and RNA was extracted from the flow-through using TRI Reagent and the Direct-zol RNA Miniprep Kit (Zymo Research). Ribosome footprints of 25–32 nt were size-selected by electrophoresis on 15% polyacrylamide TBE-Urea gels (Invitrogen), followed by rRNA depletion using the riboPOOL kit (siTOOLs Biotech). Libraries were prepared using the TruSeq Small RNA Library Preparation Kit (Illumina) and purified on 6% polyacrylamide TBE gels (Invitrogen). Single-end sequencing (50 bp, ∼400 million reads/sample) was performed on a NovaSeq X Plus instrument (Illumina).

### EpiPlex

Three replicates were performed using RNA extracted from the SH-SY5Y cell line containing the ERCC and SIRV transcript spike in mixes (E0, E1 and E2). 500ng of total RNA with RIN scores > 7 was processed using the AlidaBio EpiPlex RNA Mod Encoding Kit (AlidaBio, Product No. 100108) according to the manufacturer’s total RNA workflow. Input RNA was initially fragmented, end-polished, and ligated to sequencing adapters. The adapter-ligated RNA was then separated, with %90 of the reaction being used for enrichment of m6A and inosine (I) enrichment using the paramagnetic beads. The remaining 10% of the reaction was used as an unenriched solution control. Both the on-bead enriched fragments and the purified solution control were reverse-transcribed into cDNA, simultaneously encoding modification identity using unique modification-specific barcodes (MBCs). The resulting libraries were amplified by Index PCR to incorporate P5/P7 sequences and Unique Dual Indices (UDIs) for multiplexing. A total of 8 Cycles of PCR were performed for the enriched sample and 10 cycles of PCR were performed for the solution control sample. Prior to sequencing, the enriched and solution control libraries were quantified, pooled separately, and depleted of ribosomal RNA sequences using the BioRad SEQuoia Ribodepletion Kit (Biorad, 17006487) to generate final sequencing-ready libraries for simultaneous epitranscriptomic detection and total transcriptome analysis. EpiPlex analysis was performed using EpiScout v1.1.2 (Alida Biosciences).

### Software versions and references

*Pandas v1.5.3*

*IsoQuant v3.7.0*

*pycoQC v2.5.2*

*NumPy v1.26.4*

*SciPy v1.12.0*

*Modkit v0.3.1* [*Modkit v0.5.0* for *stats* submodule]

*Dorado v1.0.0*

*Samtools v1.18*

*Gffread v0.12.7*

*GffCompare v0.12.6*

*TD2 v1.0.6*

*GSEApy v1.1.1*

*Logomaker v0.8.7*

*GeneStructureTools v1.28.0*

*bedtools v2.30.0*

*RiboCode v1.2.11*

*gppy v0.1.4*

*pysam 0.22.1*

*biopython v1.84*

*scikit-learn v1.4.0*

*scanpy v1.9.8*

*statsmodels v0.14.1*

*CoolBox v0.4.0*

Reference genome GRCh38.p14 with GENCODE v46 annotations. The following synthetic unmodified control sequences were used: External RNA Controls Consortium (ERCC) Spike-in mix by Thermo Fisher Scientific and Spike-in RNA Variant (SIRV) by Lexogen. The reference sequences and expected titer levels were found at https://www.lexogen.com/sirvs/download/.

### DRS sequencing statistics and quantitation

Prior to alignment, base modification information stored in unaligned bam tags (*.ubam* format) was appended to *.fastq* headers using *samtools fastq (-T “*”).* Then, *IsoQuant (-d nanopore --stranded forward --complete_genedb --sqanti_output --check_canonical)* was used to align and quantify gene and transcript expression. Resultant read classifications were used to determine utilization metric. Read length statistics were based on run summaries generated by *pycoQC* ^81^.

### Gene and transcript detection and expression analysis

For gene and transcript detection, expression of greater than 1 TPM was required. For expression analysis, TPM matrices were scaled using *numpy.log1p*. To identify differential transcript usage (DTU), a permutation test was implemented that operates on transcript-level expression data grouped by gene. Only transcripts with mean expression >1 TPM and at least 3 isoforms per gene were considered. The proportional expression of transcripts within each gene was used to identify the dominant isoform (highest expressed across all replicates) and performs *scipy.stats.permutation_test* (n=1000, alternative=’greater’) comparing the dominant isoform against all other isoforms using the difference in means as the test statistic. Effect sizes were calculated as log2 fold changes between dominant and non-dominant isoforms using scaled expression values with pseudocount correction (1e-3). P values were adjusted using scipy.stats.false_discovery_control, using the default Benjamini-Hochberg procedure. To assess sequencing format effects on DTU identification, isoform expression matrices were flattened into vectors and *scipy.spatial.distance.cosine* was used to determine aggregate expression differences dependent upon sequencing format. Furthermore, *scipy.stats.entropy* was used to determine relative entropy between sequencing formats. Transcript coverage and transcript model visualizations were generated using the Python *CoolBox* package^82^.

### Transcript model and putative ORF analysis

Transcript models constructed by *IsoQuant* are stored in *.gtf* files, which were filtered based on minimum expression level (TPM > 1). Reproducible transcript models were identified using *GffCompare* and filtering for “Complete Match” (class_code = “=”). For sequencing format comparisons, an additional *GffCompare* step was performed to identify DRS-and CPS-specific models. Nucleotide sequences were generated from filtered *.gtf* files using *gffread* for subsequent ORF prediction, employing the two modules of *TD2: TD2.LongOrfs* and *TD2.Predict* (-P 0.90). ORFs with *PSAURON* score >0.9 were determined to be putative reproducibly-detected ORFs (RD-ORFs). It should be noted that TD2 can produce multiple putative ORFs from a single input transcript model. In addition, individual AD samples were processed as outlined above, excluding the replicate reproducibility requirement. Transcript model alteration classifications were determined by *IsoQuant* and outputted to *.novel_vs_known.SQANTI-like.tsv* file.

### TMT mass spectrometric validation of putative RD-ORF expression

MS/MS spectra from iNeurons study ^37^ were searched using COMET against one of four specialized concatenated target-decoy protein databases. These databases included a standard base human UniProt database (downloaded April 2024), or versions appended with custom sequences derived from DRS-derived putative RD-ORF or those from CPS data. Search parameters were set for trypsin cleavage enzyme specificity with an allowance of 1 missed cleavage event, a peptide mass tolerance of 20 ppm, and recommended fragment ion settings for low resolution MS/MS with a fragment ion bin tolerance and bin offset of 1.0005 and 0.4, respectively. Searches permitted variable modifications of methionine oxidation (+15.9949 Da) with static modifications for cysteine carbamidomethylation (+57.0215 Da) and TMT tags on lysine and peptide N termini (+304.20715 Da). Peptide spectral matches (PSMs) were filtered to a 2% FDR using the Percolator algorithm. Protein inference was subsequently performed using Parsimony and filtered to 20% FDR. Data from the ROSMAP study were searched as described above, with the following modifications. The concatenated target-decoy protein database versions were instead modified from the base UniProt database (downloaded April 2024) to contain mutually-exclusive putative RD-ORFs from Braak groupings: low (I, II), mid (III, IV), and high (V, VI). For this dataset, TMT tag modification masses on lysine and peptide N termini were set to (+229.16293) to reflect the use of TMT10 mass tags.

### Gene set enrichment and StringDB analysis

Gene sets were queried against the C5 Gene Ontology (GO) resource of Biological Processes (BP) (c5.go.bp.v2023.2.Hs) from MSigDB (www.gsea-msigdb.org/gsea/msigdb) using *gseapy.prerank*. For StringDB queries, the www.string-db.org (v12.0) web service for “Multiple Proteins” searches was used to generate interaction maps.

### Basecalling

The *Dorado* basecaller with the modification-aware RNA model (rna004\_130bps\_sup@v5.2.0) was used to independently call N^6^-methyladenosine (m6A) and adenosine-to-inosine (inosine), pseudouridine (pseU), 5-Methylcytosine (m5C) and all 2’-O-methylation (2OmeA, 2OmeC, 2OmeG, 2OmeU) RNA modifications. A reference genome containing ERCC, SIRV, and human references (hg38) and the GENCODE v46 annotation for alignment were used. The resulting *.bam* files were sorted and indexed with *Samtools.* Following basecalling, the resulting *.bam* files were divided into spike-in (ERCC and SIRV) and genomic (cell line) transcripts using *samtools view* (-L <ref.bed>) where the *.bed* files contain the chromosome information for the ERCC/SIRV genome and human genome, respectively. For cDNA base calling, the *dna_r10.4.1_e8.2_400bps_fast@v5.2.0* model was used.

### Dorado probability score distribution and modification false discovery rate (modFDR) analysis

The *modkit sample-probs* function was used to obtain a sampling distribution of probability scores for each modified base. The sample size for each *modkit sample-probs* was set at 5% of the total number of reads, with a maximum of 100,000 and a minimum of 10,042 reads. The output of this function is a histogram *.tsv* file containing the number of sampled counts for each bin of probability scores for each *.bam* file. Densities over uniform probability ranges were interpolated using *scipy.interpolate.make_smoothing_spline*(lam=1e-12). The modFDR metric was computed as the ratio of ERCC/SIRV over genomic modification score densities above a given threshold (*e.g.* P_mod_ > 0.95).

### Co-occurrence rate calculations

One million randomly sampled reads per replicate in *.bam* format were processed using *modkit extract* (--edge-filter 10, --mod-thresholds {mod_code}:0.95) to retrieve the sequence context around each modified base call. Only reads shorter than 10,000 nt were processed in order to manage memory utilization (see https://github.com/nanoporetech/modkit/issues/306). The dinucleotide co-occurrence rate was determined by first calculating the observed frequency for each dinucleotide in every replicate, and then computing the mean of these frequencies. To account for sequence biases inherent to the *Dorado* modification caller, the mean co-occurrence rates from unmodified ERCC/SIRV controls were subtracted from the rates calculated using genomic reads.

### Motif analysis of 5-mers

One million randomly sampled reads per replicate in *.bam* format were processed using *modkit extract* (--kmer-size 5, --edge-filter 10, --mod-thresholds {mod_code}:0.25) to retrieve 5-mer sequences containing a central modified base. SH-SY5Y genomic reads and ERCC/SIRV unmodified reads were separated. For each unique 5-mer sequence, a hypothesis test was used to identify significantly higher genomic modification scores compared to that of the unmodified control reads. To do this, random downsampling was first applied such that the genomic and unmodified 5-mer sets had equal depth, and then a one-sided Welch’s test was applied: *scipy.stats.ttest_ind*(equal_var=False, alternative=“greater”). P values were adjusted for multiple comparisons using *scipy.stats.false_discovery_control*(method=”bh”). To generate sequence logos, position frequency matrices were computed from the set of significant 5-mers for each modification and plotted with *logomaker.Logo*(color_scheme=”classic”).

### Compute modification percent in regions

The *UTR2UTR53* function from the *GeneStructureTools* R library was used to process UTR annotation to strand-specific UTR5 (5’-UTR) and UTR3 (3’-UTR) designations. To create region-specific bed files, *awk* was used to match “CDS”, “UTR5”, or “UTR3” features followed by sorting (*sort-k1,1-k2,2n*). Then, *bedtools subtract* (-s) was used to remove any overlapping CDS ranges from both UTR feature sets. Lastly, all bed files were sorted as before and only unique features were retained (*sort-k1,1-k2,2n | uniq*). These region-specific bed files were passed to *modkit extract* (--include-bed <region.bed>) to extract modification calls that were passed to an *awk* script to calculate read-specific modification statistics. For each read, the proportion of modified bases over the total number of corresponding base positions (both canonical and modified) was calculated. Gene-level modification percent was determined by first binning reads based on *IsoQuant* read assignment. If more than 10 reads were mapped, the modification percent for that gene was calculated as the average of the read-specific modification proportions.

### Ribo-seq processing and codon utilization

Ribo-seq reads were processed with RiboCode ^83^ to generate inferred ribosome occupancy in terms of P-site position relative to the transcriptome. Transcriptome positions were mapped to genomic positions using *gppy t2g*. Then, using *bedtools intersect*, modification position *.bedmethyl* files generated using *modkit pileup* (--edge-filter 10 --mod-thresholds {mod_code}:0.95) were intersected with genomic ribosome occupancy positions within 25 nt of the P-site. Because the Ribo-seq samples (n = 2) were generated independently of the DRS data (n = 3), all pairwise intersections were generated, resulting in six unique crosses. Sites were filtered to those with >20 Ribo-seq reads and >10 DRS reads. *Pysam* was used to extract the sequences around each site from the corresponding *.fasta* reference. Using the *biopython Seq* module, the function *translate()* was employed to convert codon sequences to corresponding amino acids. Amino acid utilization logos were generated using position frequency matrices.

### poly(A) tail length estimation and integrated features correlation

Similar to the approach described in “*Compute modification percent in regions”,* mean poly(A) tail length estimates from *Dorado* were summarized for each. Subsequently, gene-level mean poly(a) tail lengths were correlated with analogous summarized feature metrics (*e.g.* mean expression TPM or mean modification %). Specifically, all pairwise Pearson correlations used z-score-normalized and log-scaled values.

### ROSMAP TMT MS data

The Religious Orders Study and Memory and Aging Project (ROSMAP) datasets were downloaded from www.synapse.org: TMT MS (batch 5 consisting of 8 brain samples with Braak stages I, II, III, IV, IV, IV, IV, V) ^84^ and metadata ^85^.

### Integrative modeling to predict protein abundance

To assess the contribution of gene-level features to protein abundance, we performed ordinary least-squares (OLS) multivariate linear regression. All continuous variables were z-score normalized after log(x+1) transformation to place all features on a comparable scale and reduce skew: (1) RNA-seq features = TPM from DRS (mRNA); (2) Ribo-seq features = ribosome occupancy (ribo_occ); (3) DRS features = mRNA, %m6A_CDS_ (m6A_CDS) and and poly(A) tail length (polyA); (4) Combined features = mRNA, m6A_CDS, polyA, ribo_occ and m6A at the ribosomal A-site (m6A_Asite); (5) Response variable = protein abundance from MS-based quantitation. Only genes with quantified values across all relevant features were retained (inner join). The dataset was split into training (80%) and held-out test (20%) sets using a fixed random seed (random_state=1). Four nested models were fit on the training set with protein abundance as the response variable, corresponding to increasing feature complexity: (1) RNA-seq features, (2) Ribo-seq features, (3) DRS features, and (4) Combined features. Model performance was evaluated using the adjusted R², which penalizes for the number of predictors, and the global F-test p-value. Fold-change improvement in adjusted R² was calculated relative to the mRNA-only baseline model.

### Genetic algorithm-based feature selection for cell type deconvolution using m6A profiles

Cell type proportions in bulk AD brain samples were estimated using %m6A_CDS_ signature matrices derived from four iPSC-derived reference cell types: neurons (iN), astrocytes (iA), microglia (iM), and oligodendrocytes (iO). For each cell type, signature profiles were constructed by averaging replicate %m6A_CDS_ values across three replicates. Prior to feature selection, genes were pre-filtered to the top 25% most highly methylated and most variable features (by standard deviation) across the reference samples, yielding an initial candidate gene pool. A genetic algorithm (GA) was then used to identify an optimal feature subset for deconvolution. The GA was run for 250 independent trials with different random seeds using the following hyperparameters: population size = 100, maximum generations = 200, mutation rate = 0.2, tournament selection size = 5, elitism fraction = 0.25, and feature subset size constrained to 20–100 genes. These parameters established the following steps for producing each new generation of feature subsets: 1) Preserve the top 25% of genes unchanged from the previous generation (elitism). 2) Fill remaining population slots by tournament selection (tournament size = 5), in which five candidate subsets are drawn at random and the one with the lowest fitness score is chosen as a parent. This is repeated to yield 2 parent subsets. 3) Apply one-point crossover, splitting both parent gene lists at a random index and combining the first segment of one parent with the second segment of the other, with duplicate genes removed. 4) Mutate the resultant child subset (rate = 0.2) with three independent operations: adding a randomly chosen gene not already in the subset, removing a random gene from the subset, and swapping a gene for a randomly chosen alternative. This combination of elitism, tournament selection, crossover, and mutation balances exploitation of high-fitness solutions with exploration of new feature combinations.

Fitness was evaluated using a sklearn.svm.LinearSVR (C=1.0, ε=0.1, dual=True) fitted to each bulk AD sample, with non-negative coefficient clipping and proportion normalization. The fitness score penalized feature subsets that drove any cell type proportion to zero (zero-proportion penalty = 5.0 per cell type per sample) or violated the expected rank order of proportions (rank penalty = 5.0 per mis-ranked cell type), in addition to the summed absolute deviation from expected proportions derived from published single-nucleus RNA-seq data (iN=0.61, iA=0.06, iM=0.04, iO=0.28; Mathys *et al.*). Early stopping was applied if no improvement (≥0.01) was observed over 10 consecutive generations.

Genes selected across all 250 GA trials were tallied and assessed for statistical over-representation using a one-sided binomial test, where the null probability was set to the mean feature subset size divided by the initial candidate pool size. P-values were corrected for multiple testing using the Benjamini-Hochberg procedure, and genes with FDR < 0.05 were retained. The significant gene set was further refined by hierarchical clustering (*z*-score normalized), cutting the column dendrogram into six clusters and removing clusters with fewer than three members. Final cell type proportion estimates for each AD sample were obtained by fitting LinearSVR on the filtered gene set, clipping negative coefficients to zero, and normalizing weights to sum to one.

### Binomial generalized linear model (GLM) for differential m6A analysis

Differential m6A methylation across Braak stage groups was assessed using a GLM implemented in the Python *statsmodels* package. For the CDS region of each gene, the number of m6A-modified adenosines (*k*) and total adenosine observations (*n*) were extracted per sample, requiring n > 10,000 across all samples to ensure reliable rate estimates. To prevent differential sequencing depth from dominating statistical power, per-sample coverage was capped at a maximum depth of 10,000 valid adenosine sites by rescaling *k* and *n* proportionally while preserving the observed m6A fraction. The GLM modeled the binomial response [*k*, *n* - *k*] (*i.e.* m6A, unmodified A) as a function of Braak group (low: stages I–III; high: stages V–VI) with a logit link, and dispersion was estimated from the deviance (scale=“dev”), producing a binomial model that corrects standard errors and p-values for overdispersion: *statsmodels.GLM*(*k*, *n* - *k,* family=*Binomial*())*.fit*(scale=“dev”). Significance of the group coefficient (β₁, log-odds ratio) was assessed via a Wald test against a t-distribution with 11 samples − 2 degrees of freedom. Effect sizes were computed from the raw (uncapped) proportions as Δ%m6A_CDS_ and log₂ fold change (log₂(μ_high / μ_low), with a pseudocount of 10⁻⁶ to avoid division by zero). Multiple testing correction was applied using the Benjamini–Hochberg false discovery rate procedure; features with FDR < 0.05 and |log₂FC| > 0.1 were considered differentially methylated. Moreover, this GLM approach was adapted for single-nucleotide site-level differential m6A analysis, adjusting the rescaling cap to 50 valid adenosines across all candidate sites.

### AD brain single-nucleus RNA-seq (snRNA-seq) reanalysis

Single cell data from Mathys *et al.* ^65^ can be downloaded at https://compbio.mit.edu/ad_aging_brain/. Using the Python *scanpy* package, expression was depth-normalized per cell and log transformed using *scanpy.pp.normalize_total* (target_sum=1e4, layer=“counts”) and *scanpy.pp.log1p* (layer=“counts”), respectively. Using the Python *SciPy* package, the *scipy.stats.mannwhitneyu* function was used to determine differential expression between “health” and “disease” groups, followed by multiple testing correction using the Benjamini–Hochberg false discovery rate procedure.

### Use of Large Language Models (LLMs)

Enterprise implementations of Chat-GPT and Gemini were used for proofreading and for code generation. All LLM outputs were reviewed for accuracy.

## Data Availability

Sequencing data were deposited to NCBI Sequence Read Archive (SRA) under the BioProject accession PRJNA1367283.

## Code Availability

Analysis code is provided at https://github.com/danledinh/DRS_brain.

## Acknowledgments

We sincerely thank Dr. Thomas Beach and Dr. Geidy Serrano for their valuable collaborative efforts in facilitating the acquisition of brain specimens. We are grateful to the Banner Sun Health Research Institute Brain and Body Donation Program of Sun City, Arizona for the provision of human biological materials. The Brain and Body Donation Program has been supported by the National Institute of Neurological Disorders and Stroke (U24 NS072026 National Brain and Tissue Resource for Parkinson’s Disease and Related Disorders), the National Institute on Aging (P30 AG019610 and P30AG072980, Arizona Alzheimer’s Disease Center), the Arizona Department of Health Services (contract 211002, Arizona Alzheimer’s Research Center), the Arizona Biomedical Research Commission (contracts 4001, 0011, 05-901 and 1001 to the Arizona Parkinson’s Disease Consortium) and the Michael J. Fox Foundation for Parkinson’s Research. We also extend our gratitude to Tobias Bittner and Nandhini Ramamoorthi from Genentech for their efforts in sharing and coordinating the transfer of AD samples included in this study.

## Author contributions

Z. M., W. S. and D. L. conceived the study.

A. Z., X. S., A. D. S., X. C., A. N. and C. J. prepared iPSC-derived cell material.

A. B., J. L., Y. L. and W. S. prepared and sequenced libraries.

A. X-M., X. C., A. O. and Y. L. performed Ribo-seq experiments.

J. H., A. K., W. S. and D. L. performed data processing and analysis.

Z. M., W. S. and D. L. supervised the project.

L. P., M. C. and C. M. R. performed MS experiments and data analysis.

A. B., Z. M., W. S. and D. L. prepared the manuscript.

All authors discussed the results and approved the manuscript.

---

**Extended Data Figure 1.**
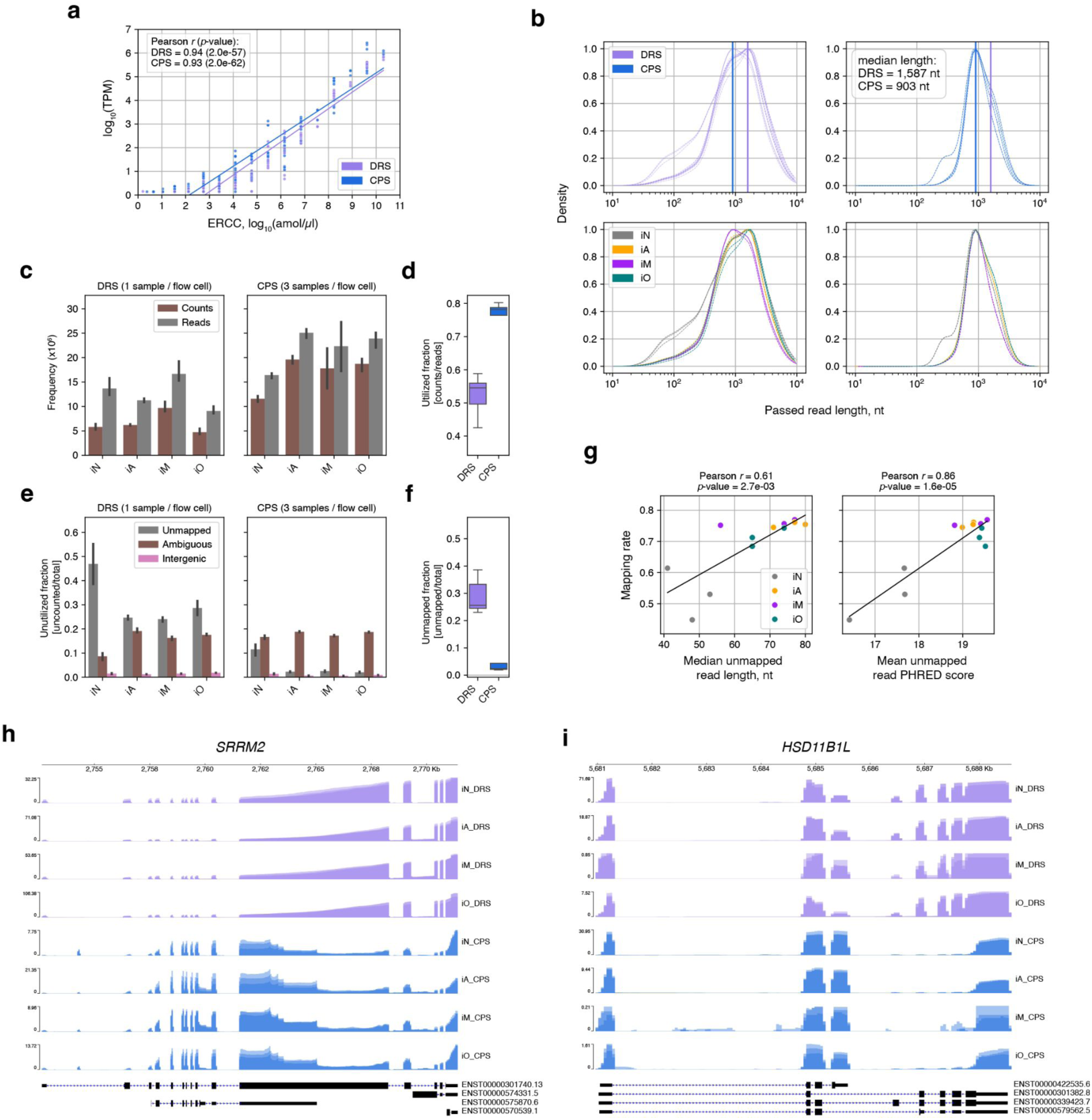
DRS versus CPS performance metrics across replicates (n = 3). **a**) Scatterplot and linear regression of observed transcript abundance versus the expected ERCC molarity, comparing between DRS and CPS. The respective Pearson correlation coefficients and corresponding P values are shown. **b**) Density plots illustrating the length distribution of reads that pass quality control filters. Density outlines in the top sub-panels are color-coded based on sequencing format: DRS versus CPS. The vertical lines indicate respective median read length along the density distribution. The median read length values are also shown. Density outlines in the bottom sub-panels are color-coded based on sample type: iNeuron (iN), iAstrocyte (iA), iMicroglia (iM) and iOligodendrocyte (iO). **c**) Barplots displaying the mean read and count yields per sample, with error bars indicating the 95% confidence interval, for DRS and for CPS. Counts are defined as reads that are utilized to measure gene expression. **d**) Boxplots showing the utilization fraction for DRS and for CPS, which is defined as counts divided by total reads. **e**) Barplots displaying the mean utilization fraction per sample, disaggregated by unutilized categories: unmapped, ambiguous and intergenic. The error bars indicate the 95% confidence interval for DRS and for CPS. **f**) Boxplot depicting the unmapped fraction for DRS and for CPS. **g**) Scatterplot and linear regression of DRS total library mapping rate as a function of unmapped read features: median read length and of mean read PHRED score. Each point represents individual samples and are color-coded based on cell type: iNeuron (iN), iAstrocyte (iA), iMicroglia (iM) and iOligodendrocyte (iO). Respective regression lines, Pearson correlation coefficients and P values are shown. **h**) For the SRRM2 gene, Integrated Genome Viewer (IGV) coverage tracks. Tracks are color-coded by sequencing format: DRS and CPS. **I**) Similar to before, except for the HSD11B1L.

**Extended Data Figure 2.**
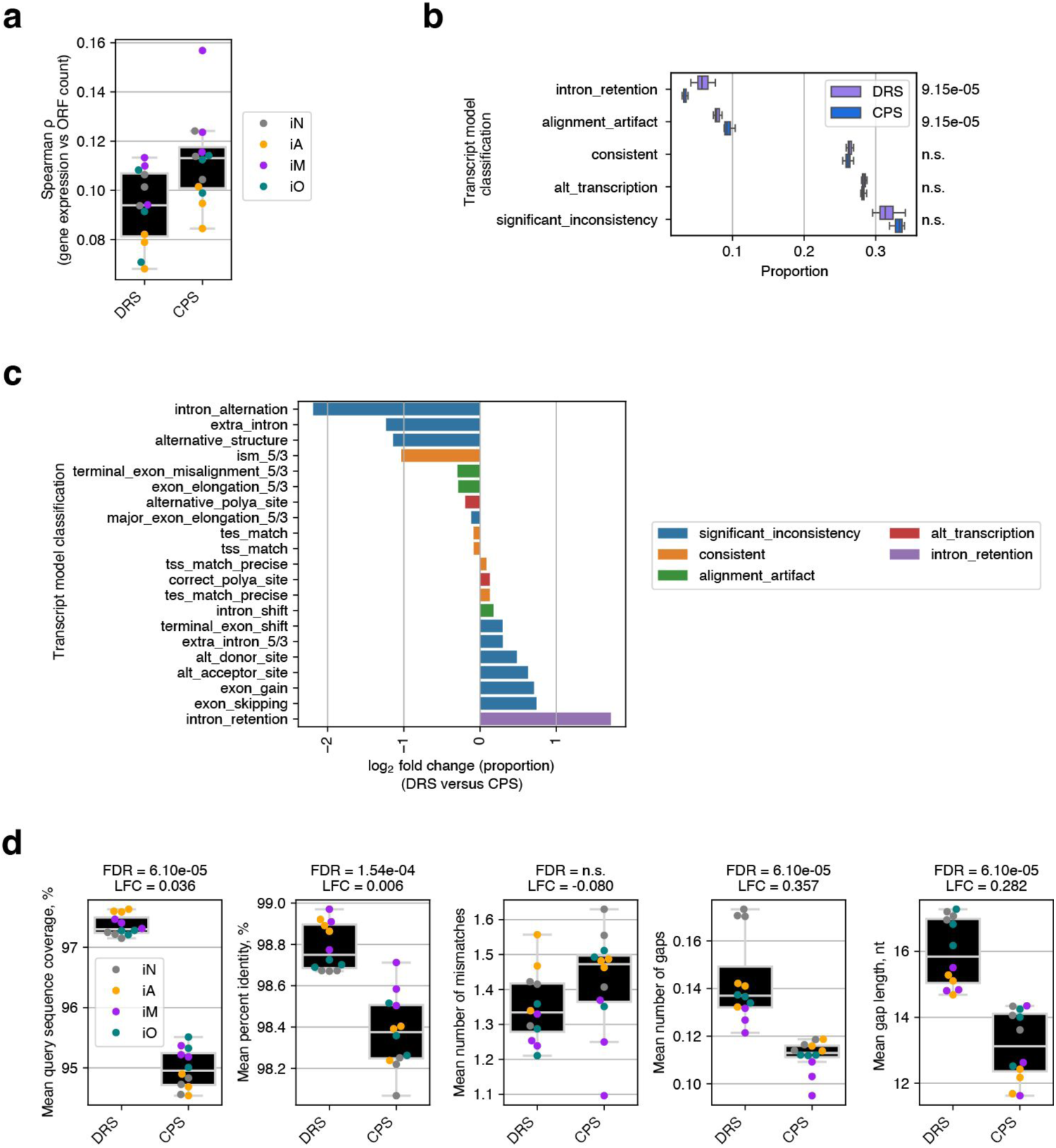
Transcript model and ORF analysis. **a**) Boxplot depicting Spearman correlation ⍴ values for the association between gene expression and observed ORF counts for each sample across DRS and CPS formats, with color-coding by cell type. n= 3 per cell type. **b**) Boxplot showing high-level transcript model classification proportions across DRS and CPS formats, color-coded by format. Mann-Whitney *U*-test FDR values shown on the right y-axis. **c**) Barplot showing significant log2 fold change in detailed transcript model classification proportions, contrasting DRS versus CPS. Bars are colored by high-level classification. **d**) Boxplots showing alignment metrics from putative ORF query of RefSeq database using *BLASTp*, across DRS and CPS formats and color-coded by iPCS-derived cell type. From left to right: mean query sequence coverage (%), mean percent identity (%), mean number of mismatches, mean number of gaps, and mean gap length (nt). Mann-Whitney *U*-test FDR values and log_2_ fold change (LFC) are shown.

**Extended Data Figure 3.**
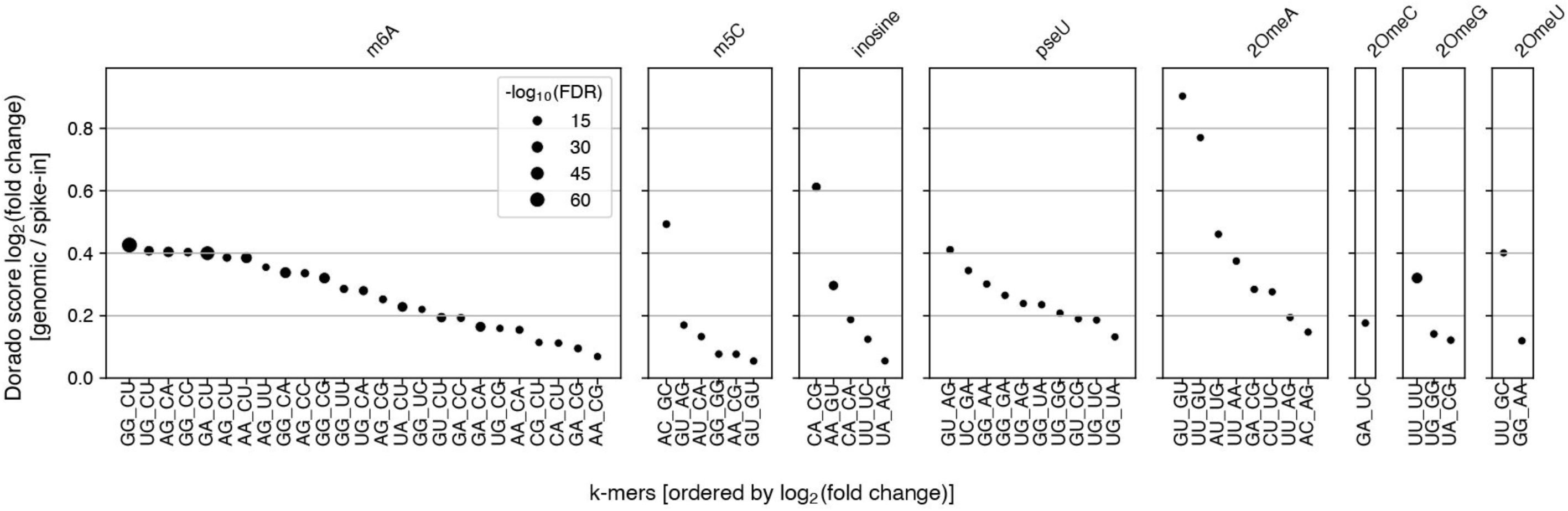
5-mer sequences with significantly higher RNA modification probability. Scatter plots showing 5-mers with significantly higher RNA modification probabilities in the genomic context compared to an unmodified synthetic random sample, across replicates (n = 3). The 5-mers are ranked based on the magnitude of modification probability fold change. The central base denoted as “_” is the site of modification. A one-sided Welch’s *t*-test together with Benjamini-Hochberg multiple testing P value adjustment was used to determine significance. Point sizes are scaled to the negative log_10_ transform of FDR.

**Extended Data Figure 4.**
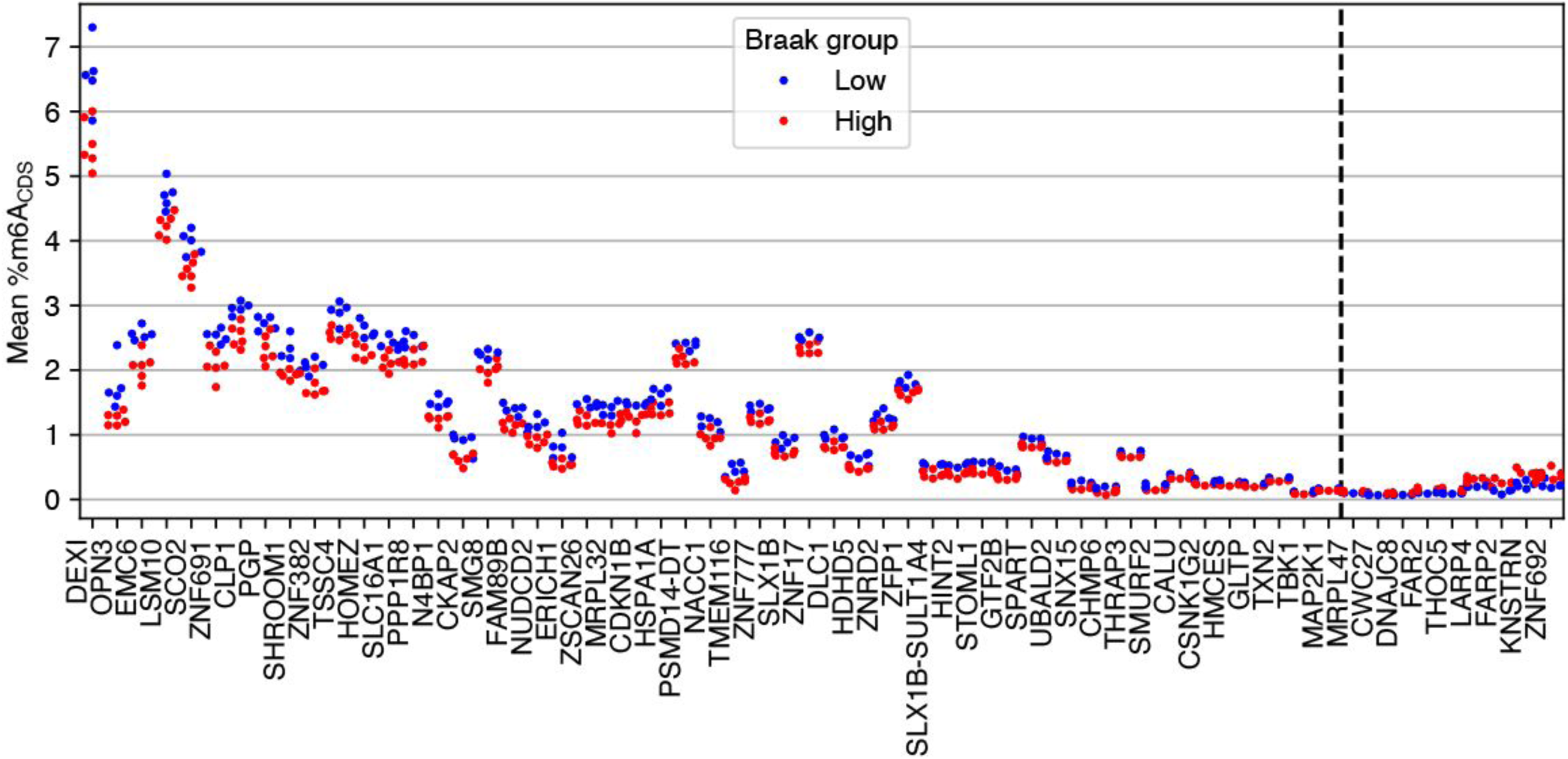
Swarmplot showing differentially methylated genes by gene-level mean %m6A_CDS_, color-coded by Braak group (low versus high). Genes sorted by absolute difference in mean %m6A_CDS_.

**Supplemental Table 1.**
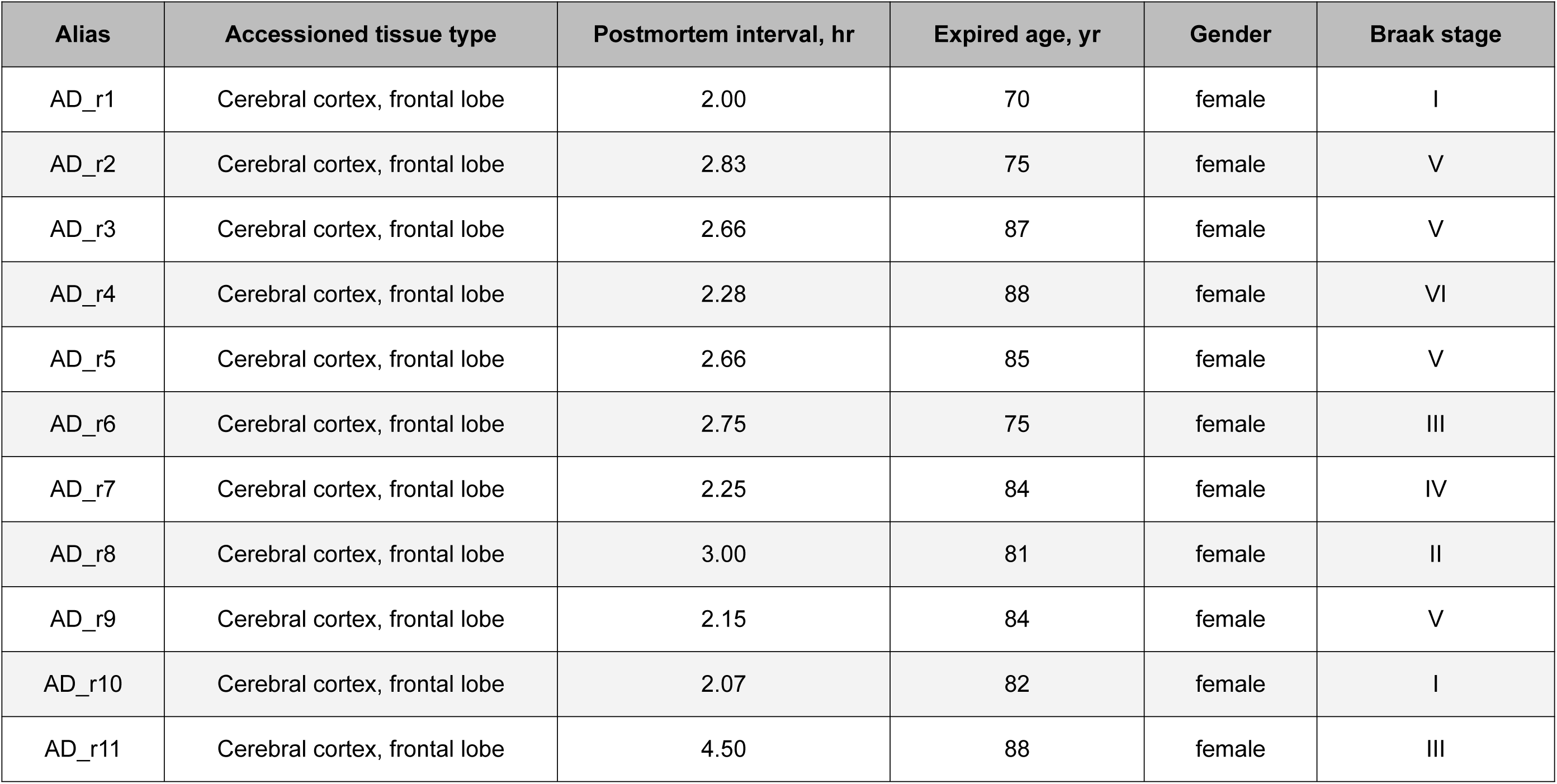

| **Alias** | **Accessioned tissue type** | **Postmortem interval, hr** | **Expired age, yr** | **Gender** | **Braak stage** |
| --- | --- | --- | --- | --- | --- |
| AD_r1 | Cerebral cortex, frontal lobe | 2.00 | 70 | female | I |
| AD_r2 | Cerebral cortex, frontal lobe | 2.83 | 75 | female | V |
| AD_r3 | Cerebral cortex, frontal lobe | 2.66 | 87 | female | V |
| AD_r4 | Cerebral cortex, frontal lobe | 2.28 | 88 | female | VI |
| AD_r5 | Cerebral cortex, frontal lobe | 2.66 | 85 | female | V |
| AD_r6 | Cerebral cortex, frontal lobe | 2.75 | 75 | female | III |
| AD_r7 | Cerebral cortex, frontal lobe | 2.25 | 84 | female | IV |
| AD_r8 | Cerebral cortex, frontal lobe | 3.00 | 81 | female | II |
| AD_r9 | Cerebral cortex, frontal lobe | 2.15 | 84 | female | V |
| AD_r10 | Cerebral cortex, frontal lobe | 2.07 | 82 | female | I |
| AD_r11 | Cerebral cortex, frontal lobe | 4.50 | 88 | female | III |

